# Continuous-trait probabilistic model for comparing multi-species functional genomic data

**DOI:** 10.1101/283093

**Authors:** Yang Yang, Quanquan Gu, Yang Zhang, Takayo Sasaki, Julianna Crivello, Rachel J. O’Neill, David M. Gilbert, Jian Ma

## Abstract

A large amount of multi-species functional genomic data from high-throughput assays are becoming available to help understand the molecular mechanisms for phenotypic diversity across species. However, continuous-trait probabilistic models, which are key to such comparative analysis, remain underexplored. Here we develop a new model, called phylogenetic hidden Markov Gaussian processes (Phylo-HMGP), to simultaneously infer heterogeneous evolutionary states of functional genomic features in a genome-wide manner. Both simulation studies and real data application demonstrate the effectiveness of Phylo-HMGP. Importantly, we applied Phylo-HMGP to analyze a new cross-species DNA replication timing (RT) dataset from the same cell type in five primate species (human, chimpanzee, orangutan, gibbon, and green monkey). We demonstrate that our Phylo-HMGP model enables discovery of genomic regions with distinct evolutionary patterns of RT. Our method provides a generic framework for comparative analysis of multi-species continuous functional genomic signals to help reveal regions with conserved or lineage-specific regulatory roles.

## Introduction

Multi-species functional genomic data from various high-throughput assays (e.g., ChIP-seq of transcription factor proteins or histone marks) are highly informative for the comparative analysis of gene regulation conservation and differences between human and other mammalian species (Villar et al., 2015; Cotney et al., 2013; Brawand et al., 2011; Levin et al., 2016). Continuous-trait models, which are key to the modeling of functional genomic signals, are gaining increasing attention in genome-wide comparative genomic studies (Naval-Sánchez et al., 2015; Rohlfs et al., 2013). However, computational models are under-explored to fully model continuous functional genomic data in the context of multi-species comparisons. In particular, to the best of our knowledge, there are no existing algorithms available to simultaneously infer heterogeneous continuous-trait evolutionary models along the entire genome.

Several types of continuous-trait evolutionary models have been developed for individual loci. One basic model (Felsenstein, 1985; Pagel, 1999; Freckleton, 2012) assumes that continuous traits evolve by Brownian motion. This model has been extended to more complicated Gaussian processes such as the Ornstein-Uhlenbeck (OU) process (Hansen, 1997; Butler and King, 2004; Hansen et al., 2008). However, the existing methods that use continuous-trait evolutionary models in comparative genomics either apply a single evolutionary model to signals of selected regions, or test different evolutionary model assumptions with prior knowledge at selected regions (Rohlfs et al., 2013; Brawand et al., 2011; Naval-Sánchez et al., 2015; Drummond et al., 2012). In other words, the continuous-trait evolutionary models have not been utilized in simultaneously estimating heterogeneous phylogenetic trees across different loci along the entire genome based on functional genomic data.

In this paper, we develop a new continuous-trait probabilistic model for more accurate state estimation based on multivariate features from different species using functional genomic signals. We call our model phylogenetic hidden Markov Gaussian processes (Phylo-HMGP). Our new method incorporates the evolutionary affinity among multiple species into the hidden Markov model (HMM) for exploiting both temporal dependencies across species in the context of evolution and spatial dependencies along the genome in a continuous-trait model. Note that our Phylo-HMGP is fundamentally different from the existing models that are restricted to discrete state space of the studied traits (Siepel and Haussler, 2005; Hobolth et al., 2007; Liu et al., 2014; Jensen and Pedersen, 2000; Lunter and Hein, 2004; Qu et al., 2018). In particular, Phylo-HMMs define a stochastic process of discrete-trait character changes (Siepel and Haussler, 2005), where different states estimated by Phylo-HMMs can reflect different patterns of substitutions or background distributions. However, Phylo-HMMs do not handle continuous signals. The models in (Jensen and Pedersen, 2000; Lunter and Hein, 2004) share similar mechanisms and have the same limitations. In the recent phylo-epigenetic model (Qu et al., 2018), the nature of the method is still based on transitions between discrete levels of observed signals, with the need to discretize the traits. In this work, our Phylo-HMGP explores a new integrated attempt to utilize continuous-trait evolutionary models with spatial constraints to more effectively study the genome-wide features across species. Our model is also flexible such that various continuous-trait evolutionary models or assumptions can be incorporated according to the actual problems. We believe that Phylo-HMGP provides a generic framework, which can be applied to different types of functional genomic signals, to more precisely capture the evolutionary history of regulatory regions across different species.

In this work, we generated a new cross-species DNA replication timing (RT) dataset from the same cell type in five primate species (human, chimpanzee, orangutan, gibbon, and green monkey). The RT program in eukaryotic cells duplicates the genome with a highly regulated temporal pattern. Genome-wide RT maps have revealed replication domains that correlate with higher order genome organization such as Hi-C A/B compartments and topologically associating domains (TADs) (Rhind and Gilbert, 2013; Ryba et al., 2010; Pope et al., 2014; Dileep et al., 2015; Solovei et al., 2016). It is known that RT changes across half of the genome during cell differentiation and disease (Ryba et al., 2011, 2012; Yue et al., 2014; Rivera-Mulia et al., 2015; Dileep et al., 2015). In addition, studies have shown conservation of RT between human and mouse (Yaffe et al., 2010; Ryba et al., 2010; Yue et al., 2014; Pope et al., 2014). Microscopy also revealed that chromosome regions with early and late RT have specific spatial localization preferences in nucleus that are conserved in evolution (Solovei et al., 2016). However, we have limited understanding of how the RT program has evolved in mammals. To the best of our knowledge, there is no existing study to investigate the RT conservation and dynamics for more than two mammalian species beyond the human-mouse comparison. Here we apply Phylo-HMGP to reveal genome-wide distributions of distinct evolutionary patterns of RT in five primates. We found that constitutive early and constitutive late RT regions, as defined from human ES cell differentiation (Ryba et al., 2011; Dileep et al., 2015), exhibit a strong correlation with the predicted conserved early RT and conserved late RT patterns. We also found distinct gene functions associated with different RT evolution patterns. In addition, the predicted RT patterns across species show correlations with other genomic and epigenomic features, including higher order genome organization, *cis*-regulatory elements, chromatin marks, and transposable elements. Our results from the comparative RT analysis in five primate species demonstrate the potential of our Phylo-HMGP model to help reveal regions with conserved or lineage-specific regulatory roles for the entire genome.

## Results

### Overview of the phylogenetic hidden Markov Gaussian processes (Phylo-HMGP) model

Here we first provide an overview of the proposed model (Fig. 1). The details of the model are described in the Methods section. Our model aims to estimate different evolutionary patterns from multi-species functional genomic signals. As illustrated in Fig. 1C, the input contains the observed continuous-trait signals from orthologous genomic regions from multiple species. The output is a genome-wide partition where neighboring genomic segments have different predicted states of multi-species signals, reflecting different evolution patterns of the signals being considered.

**Figure 1:**
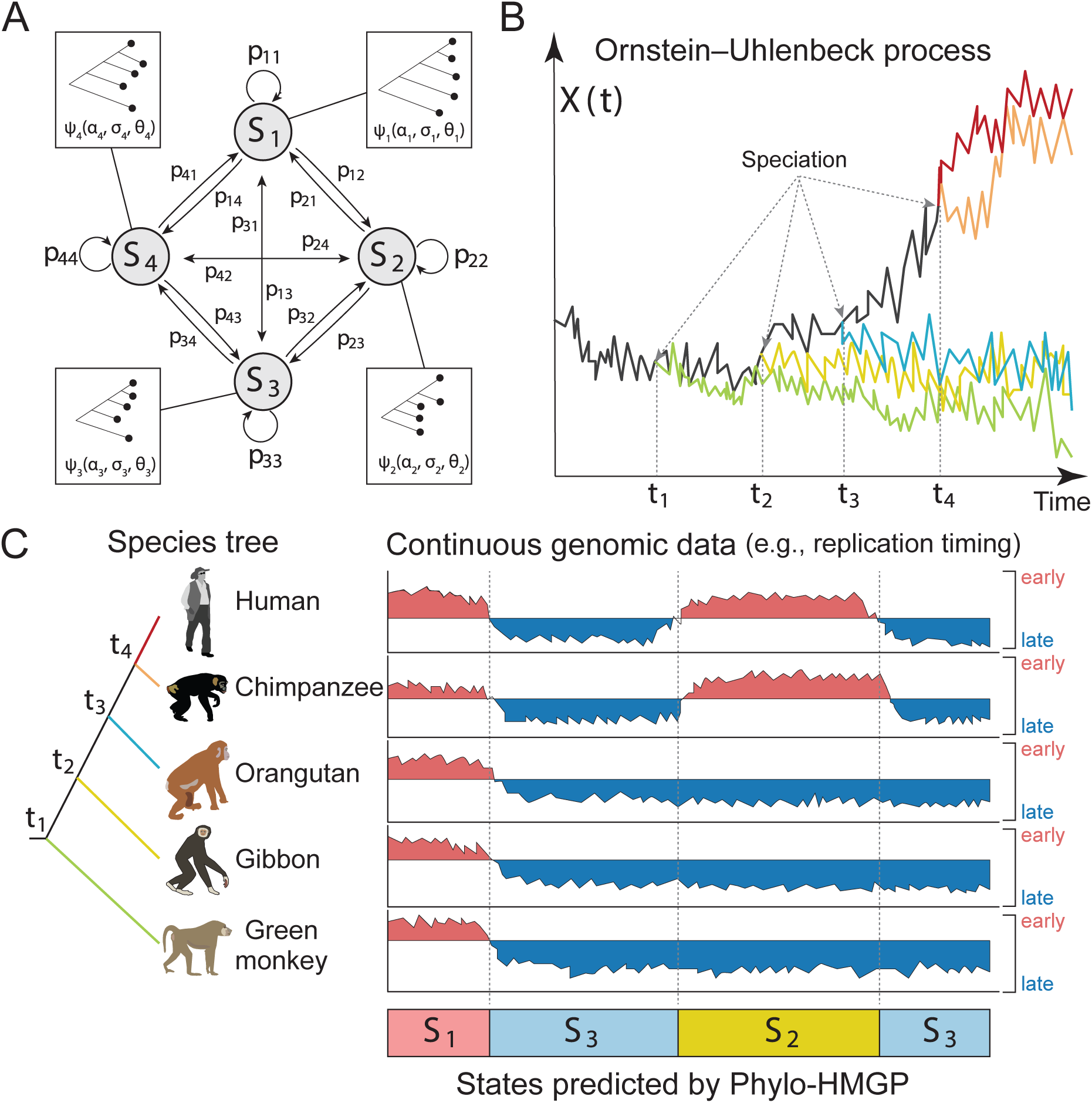
Overview of the Phylo-HMGP model. **(A)** Example of the state space and state transition probabilities of the Phylo-HMGP model associated with the continuous genomic data in **(C)**. *S*_*i*_ represents a hidden state. Each hidden state is determined by a phylogenetic model *ψ*_*i*_, which is parameterized by the selective strengths *α*_*i*_, Brownian motion intensities *σ*_*i*_, and the optimal values *θ*_*i*_ of ancestor species and observed species on the corresponding phylogenetic tree. *α*_*i*_, *σ*_*i*_, and *θ*_*i*_ are all vectors. **(B)** Illustration of the Ornstein-Uhlenbeck (OU) processes along the species tree specified in **(C)**. *X*(*t*) represents the continuous trait at time *t*. The trajectories of different colors along time correspond to the evolution of the continuous trait in different lineages specified by the corresponding colors in **(C)**, respectively. The time points *t*_1_, *t*_2_, *t*_3_, and *t*_4_ represent the speciation time points, which correspond to the speciation events shown in **(C)**. The observations of the five species also represent an example of state *S*_2_ in **(C). (C)** Simplified representation of input and output of the Phylo-HMGP model. The five tracks of continuous signals represent the observation from five species. *S*_*i*_ represents the underlying hidden states. Specifically, the example is the replication timing data, where ‘early’ and ‘late’ represent the early and late stages of replication timing, respectively. The species tree alongside the continuous data tracks shows the evolutionary relationships among the five species in this study. See also Fig. S2, Fig. S8, and Fig. S9.

We define a Phylo-HMGP model as **h** = (*S, ψ, A, π*), where *S* is the set of states, *ψ* is the set of phylogenetic models, *A* is the state-transition probability matrix, and *π* represents the initial state probabilities, respectively. Suppose there are *M* hidden states. We have *S* = {*s*_1_, …, *s*_*M*_}, *ψ* = {*ψ*_1_, …, *ψ*_*M*_}, *A* = {*a*_*ij*_}, 1 ≤*i, j* ≤*M*, and *π* = {*π*_1_, …, *π*_*M*_}. Fig. 1A shows the state space where different states are associated with varied phylogenetic tree models. Each phylogenetic tree model is parameterized with the OU processes, an example of which is shown in Fig. 1B. Note that in this paper, we focus on the Ornstein-Uhlenbeck (OU) process and apply it to analyze cross-species RT data. We also discuss and compare with Brownian motion process within the framework (see later section), which is also used as the Gaussian process for realizations of *ψ*_*j*_ to construct the emission probability distributions, *j* = 1, …, *M*. *ψ*_*j*_ differs under different evolutionary models. Other Gaussian processes can also be embedded into the framework by alternative definitions of *ψ*_*j*_.

Phylo-HMGP provides a generic framework to more effectively incorporate multi-species functional genomic data into the HMM for analyzing both temporal dependencies across species in the phylogeny and spatial dependencies along the entire genome in a continuous-trait model. The framework is flexible and the Gaussian processes embedded in the HMM can be specialized to different evolutionary models (e.g., in this work we focus on OU processes to construct the phylogenetic tree model). The source code of Phylo-HMGP can be accessed at: https://github.com/ma-compbio/Phylo-HMGP.

### Simulation study demonstrates the robustness of Phylo-HMGP

To explore whether incorporating evolutionary temporal constraints into the HMM can improve the accuracy of identifying different evolutionary patterns, we applied our method to 12 synthetic datasets in two types of simulation studies. We assessed the performance based on Adjusted Mutual Information (AMI), Normalized Mutual Information (NMI), Adjusted Rand Index (ARI), Precision, Recall, and *F*_1_ score (Manning et al., 2008; Vinh et al., 2010) by comparing the predicated states with the ground truth states (Supplementary Methods A.6). We used HMM to generate the samples in simulation study I (SS-I), while simulation study II (SS-II) did not use HMM and was instead based on a Gaussian Mixture Model (GMM). Both SS-I and SS-II contained six synthetic datasets (sample size = 50,000 each), respectively. Detailed descriptions of the simulated datasets are in the Methods section.

We compared Phylo-HMGP-OU and Phylo-HMGP-BM with the Gaussian-HMM method, the GMM method, and the K-means clustering method in both SS-I and SS-II. For each dataset, we ran each method 10 times. Each method was started from different initializations and given the state number as 10. We reported the average performance of the 10 runs as the final performance of the respective method. We applied the same regularization parameter to Phylo-HMGP-OU on all of the 12 datasets, without tuning the parameter specifically on each dataset. The results show that Phylo-HMGP-OU outperforms the other methods on AMI, ARI, and *F*_1_ score on all of the six datasets in SS-I (Fig. 2A and Table. S1). In particular, Phylo-HMGP-OU shows significant advantage in reaching higher ARI on average in different datasets, as compared to the other methods. In SS-II (Fig. 2B and Table. S2), the performance of Phylo-HMGP-OU decreases occasionally (SS-II-1 and II-2) as compared to its performance in SS-I. However, Phylo-HMGP-OU still outperforms the other methods in five of the six datasets. Phylo-HMGP-BM reaches the highest performance on SS-I-1, while Phylo-HMGP-OU maintains comparable performance to Phylo-HMGP-BM on this dataset. These simulation results strongly suggest that Phylo-HMGP-OU can achieve robust performance even when the data are simulated from a non-HMM model such as the Gaussian mixture model. Note that in the rest of the Results section, we use “Phylo-HMGP” to refer to “Phylo-HMGP-OU”.

**Figure 2:**
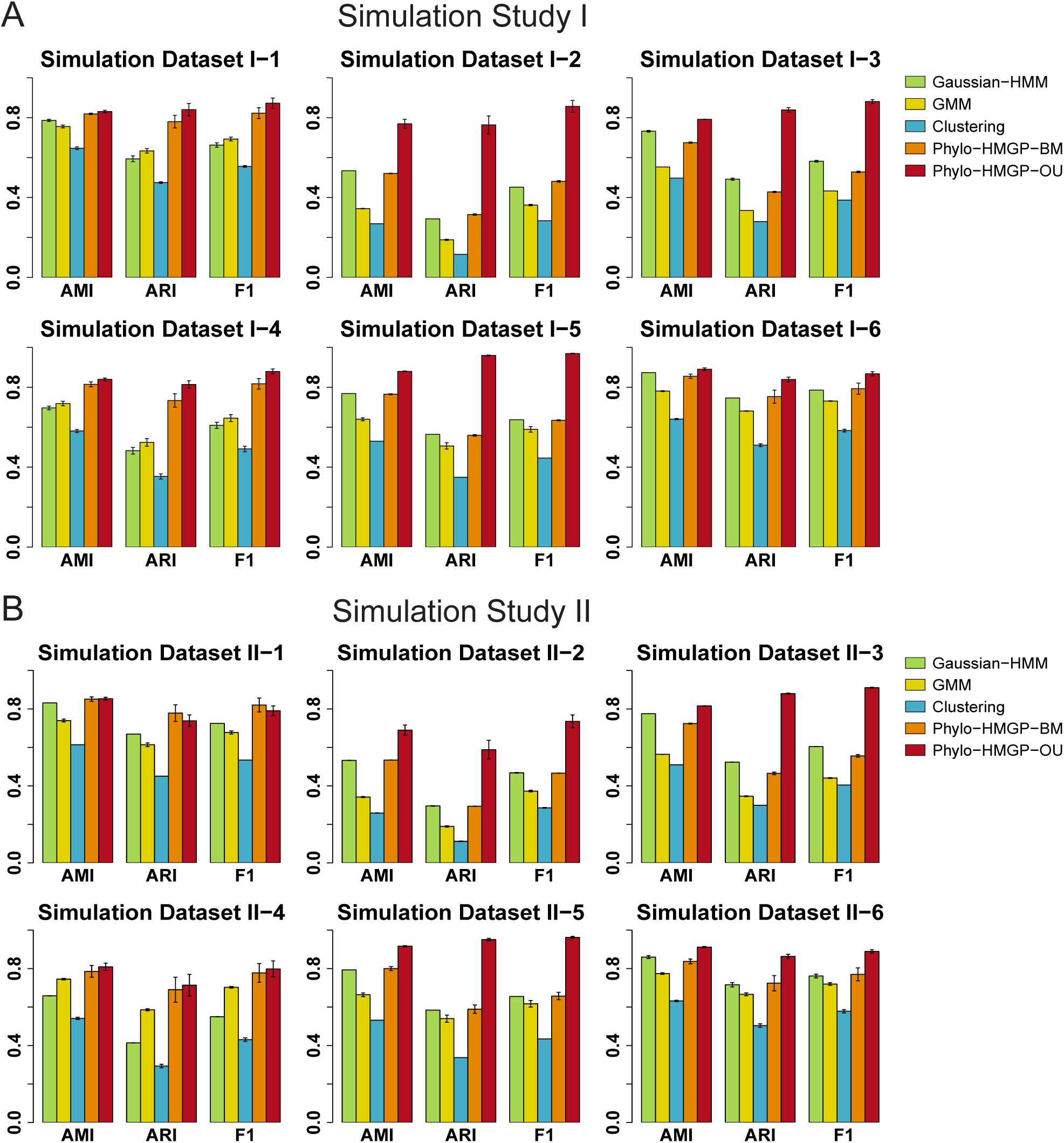
Evaluation using simulated datasets. **(A)** Evaluation of Gaussian-HMM, GMM, K-means Clustering, Phylo-HMGP-BM, and Phylo-HMGP-OU on six simulation datasets in Simulation Study I in terms of AMI (Adjusted Mutual Information), ARI (Adjusted Rand Index), and *F*_1_ score. **(B)** Evaluation of Gaussian-HMM, GMM, K-means Clustering, Phylo-HMGP-BM, and Phylo-HMGP-OU on six simulation datasets in Simulation Study II in terms of AMI, ARI, and *F*_1_ score. In both **(A)** and **(B)**, the standard error of the results of 10 repeated runs for each method is also shown as the error bar. See also Table S1, Table S2, and Fig. S1.

### Phylo-HMGP reveals genome-wide patterns of replication timing across primate species

Next, we applied the Phylo-HMGP method to study different evolutionary patterns of RT in mammalian genomes. We generated genome-wide RT maps based on Repli-seq (Marchal et al., 2017) in lymphoblastoid cells from five primate species, including human, chimpanzee, orangutan, gibbon, and green monkey. See Methods section for the details on how we processed the data. We then applied Phylo-HMGP to this multi-species RT dataset. We set the state number as 30 based on estimation from K-means clustering (see Supplementary Methods A.8 and Fig. S3). We identified both conserved and lineage-specific states with differences in RT patterns across species. Here we classified the 30 states into five groups: conserved early (denoted as E), conserved late (L), weakly conserved early (WE), weakly conserved late (WL), and non-conserved (NC) (Supplementary Methods A.9). In the E group, all five species have early RT. In the WE group, four species have early RT. We assign states to the L group and the WL group similarly. The remaining states are assigned to the NC group.

The representative RT signal patterns of the 30 predicted states are shown in Fig. 3A, with examples of the states and groups shown in Fig. 3B and Fig. 3D. Distributions of RT signals of the five species in each of the 30 states are shown in Fig. S2, including other lineage-specific patterns, conserved patterns, or divergent patterns. States 1-8 are conserved early or conserved late states of RT, making up approximately 47.7% of the whole genome. States 9-18 display different lineage-specific RT patterns. State 9 (Fig. 3B) and 10 represent human-chimpanzee (hominini) specific patterns of early RT and late RT, respectively. State 11 shows human-chimp-orangutan (hominid) specific early RT. States 12-18 reflect single-lineage specific patterns, where one species differs from all the other species.

**Figure 3:**
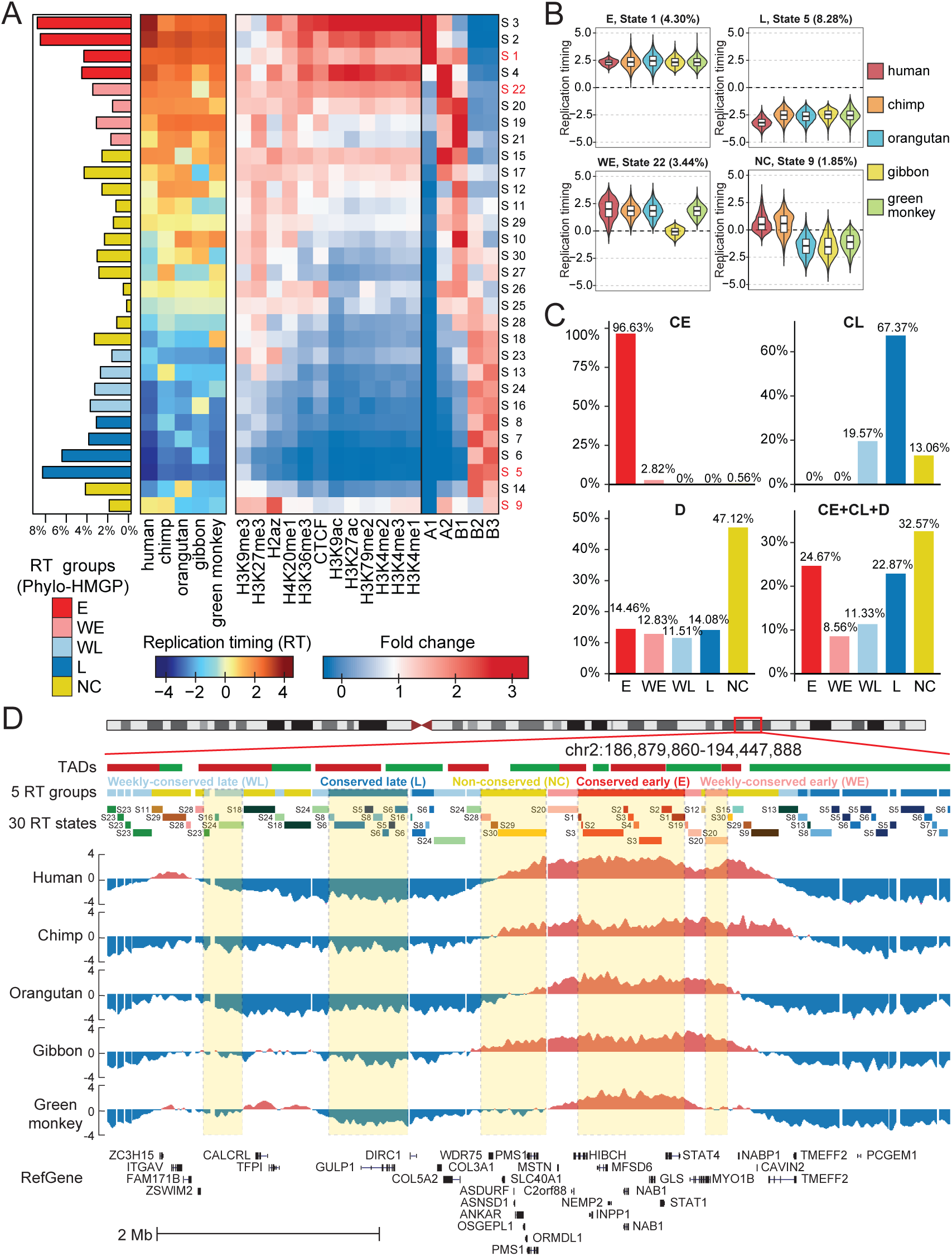
RT evolution patterns identified by Phylo-HMGP. **(A)** Panel 1 (leftmost): Proportions of the 30 RT states on the entire genome. The RT states are categorized into 5 groups: conserved early (E), weakly conserved early (WE), weakly conserved late (WL), conserved late (L), and other stages (NC), respectively. Panel 2: Patterns of the 30 states. Each row of the matrix corresponds to the state at the same row in Panel 1, and columns are species. Each entry represents the median of the RT signals of the corresponding species in the associated state. Panel 3: Enrichment of different types of histone marks and CTCF binding site (higher fold change represents higher enrichment). Panel 4: Enrichment of subcompartment A1, A2, B1, B2, and B3. **(B)** Four examples of RT signal distributions in states with different patterns (State 1: E; State 5: L; State 22: WE; State 9: NC with human-chimpanzee specific early RT). **(C)** Comparison of predicted RT patterns with the constitutively early/late RT regions identified across cell types. **(D)** Examples of different RT states and RT groups in five species predicted by Phylo-HMGP. TADs called by the Directionality Index method are shown at the top. See also Fig. S2, Fig. S3, Fig. S4, Fig. S5, Fig. S6, and Fig. S7.

Phylo-HMGP estimated the transition probabilities between the 30 predicted states (Fig. S4). We noticed that overall the transition probabilities are higher within the E group and L group. Phylo-HMGP also simultaneously estimated the model parameters of selection strength, Brownian motion intensity, and optimal values of the phylogenetic model associated with each state (Fig. S5 and Fig. S6). We found that the estimated parameters correspond very well to the lineage-specific RT patterns. For example, for state 9 and 10, the human-chimpanzee specific states, the estimated strongest Brownian motion intensity happened on the branch leading to human and chimpanzee, and strong selection strength is also estimated for human and chimpanzee. We observed similar correlations for other states. We also compared Phylo-HMGP with the other methods on an evaluation dataset constructed from the RT data and found that Phylo-HMGP outperforms other methods (Supplementary Results B.1 and Fig. S7).

### RT evolution patterns correlate with A/B compartments and histone marks

Analysis based on Hi-C data has shown that the genome can be divided into two compartments called A/B compartments (Lieberman-Aiden et al., 2009), with at least five subcompartments, namely A1, A2, B1, B2 and B3, which have different genomic and epigenomic properties (Rao et al., 2014). A1 and A2 subcompartments both show early RT, with the difference that replications in A2 regions finish later than A1. B2 and B3 subcompartments show late RT, while replications in B1 happen in the middle of S-phase (Rao et al., 2014). We used the subcompartment definitions in the human lymphoblastoid cell line GM12878 from (Rao et al., 2014) and calculated the enrichment of the five subcompartments in the 30 predicted RT states. We observed that different predicted RT evolution patterns show distinct enrichments of the subcompartments. For example, the predicted RT states in the E group (state 1-4) show the strongest correlation with A1 or A2, while the predicted RT states in the L group are enriched with B2 and B3. The majority of the states in the NC group are most enriched with A2 or B1. States in the WE group and WL group are enriched with A2/B1, and B2/B3, respectively.

We next compared the enrichments of different histone marks and CTCF binding site within each RT state. Fig. 3A panel 3 shows the enrichment distributions of histone marks and CTCF binding site across the five predicted RT groups. These distributions are consistent with the epigenomic feature patterns of the subcompartments that are enriched in the corresponding states. We found that RT states in the E group show strong positive correlation with active histone marks (e.g., H3K27ac, H3K36me3, and H3K4me1) and the CTCF binding sites. On the contrary, RT states in the L/WL groups show distinct depletion of these histone marks and the CTCF binding sites. The majority of predicted states in the NC group instead exhibit variations in the enrichments of different types of histone marks.

Among the NC states, state 9 is identified as a human-chimpanzee specific early RT state (Fig. 3B). It displays a unique pattern of histone mark enrichment, showing the strongest correlation with H2A.Z (*p*-values<1e-07) as compared to other predicted states. Recent studies have reported that H2A.Z is progressively enriched towards early RT loci (Du et al., 2018). Another state with interesting features is state 4, a conserved early state. The RT is significantly early in human in state 4, similar to other states in the E group. All of the other states in the E group (state 1-3) are strongly correlated with A1 subcompartment. State 4, however, is enriched with A2 subcompartment and is more positively correlated with H3K9me3, which generally has stronger enrichment in A2 than A1 (Rao et al., 2014). Therefore, state 4 represents a distinct state in the E group. These results demonstrate that Phylo-HMGP has the sensitivity to distinguish within similar evolutionary patterns of RT.

### Different RT evolution patterns reflect different functions

Previous studies have shown that different genomic regions have different levels of cell type specificity for RT, including constitutively early, constitutively late, and more dynamic across different cell types (Ryba et al., 2011; Dileep et al., 2015). We compared the states from Phylo-HMGP with the constitutive and developmental RT patterns discovered during ES cell differentiation (Dileep et al., 2015), including constitutively early (CE), constitutively late (CL), developmentally regulated (D), and undetermined. We found that overall the constitutively early or constitutively late RT regions in the human genome have high consistency with the strongly conserved RT evolution patterns (Fig. 3C). The findings are consistent with previous observations in human-mouse RT comparison (Ryba et al., 2011).

Among the CE regions that are also covered in the cross-species RT comparisons by Phylo-HMGP, 99.45% of the regions are assigned to the states of conserved early or weakly conserved early (*p*-value<2.2e-16). Also, 86.94% of the CL regions in human are within states of conserved late or weakly conserved late (*p*-value<2.2e-16). In contrast, the D regions show more diverse patterns across the five RT groups predicted by Phylo-HMGP. This also suggests that the RT regions in lymphoblastoid cells with similar RT profile across different cell types are highly likely to be conserved in primates. However, a significant fraction of conserved RT regions in primates also shows cell-type specific RT patterns in human. We performed Gene Ontology (GO) analysis for the conserved RT early regions with respect to the constitutive/non-constitutive RT patterns using DAVID (Huang et al., 2007), and found clear differences in gene functions (Table S3). We further performed GO analysis for the lineage-specific RT states (see Fig. 4A and Table S4). We found that genes associated with different states have different functions and biological processes. For example, the hominini-specific early RT state (state 9) is enriched with genes having sensory functions. These analyses suggest that regions with different RT evolution patterns may contain genes with distinct functions.

**Figure 4:**
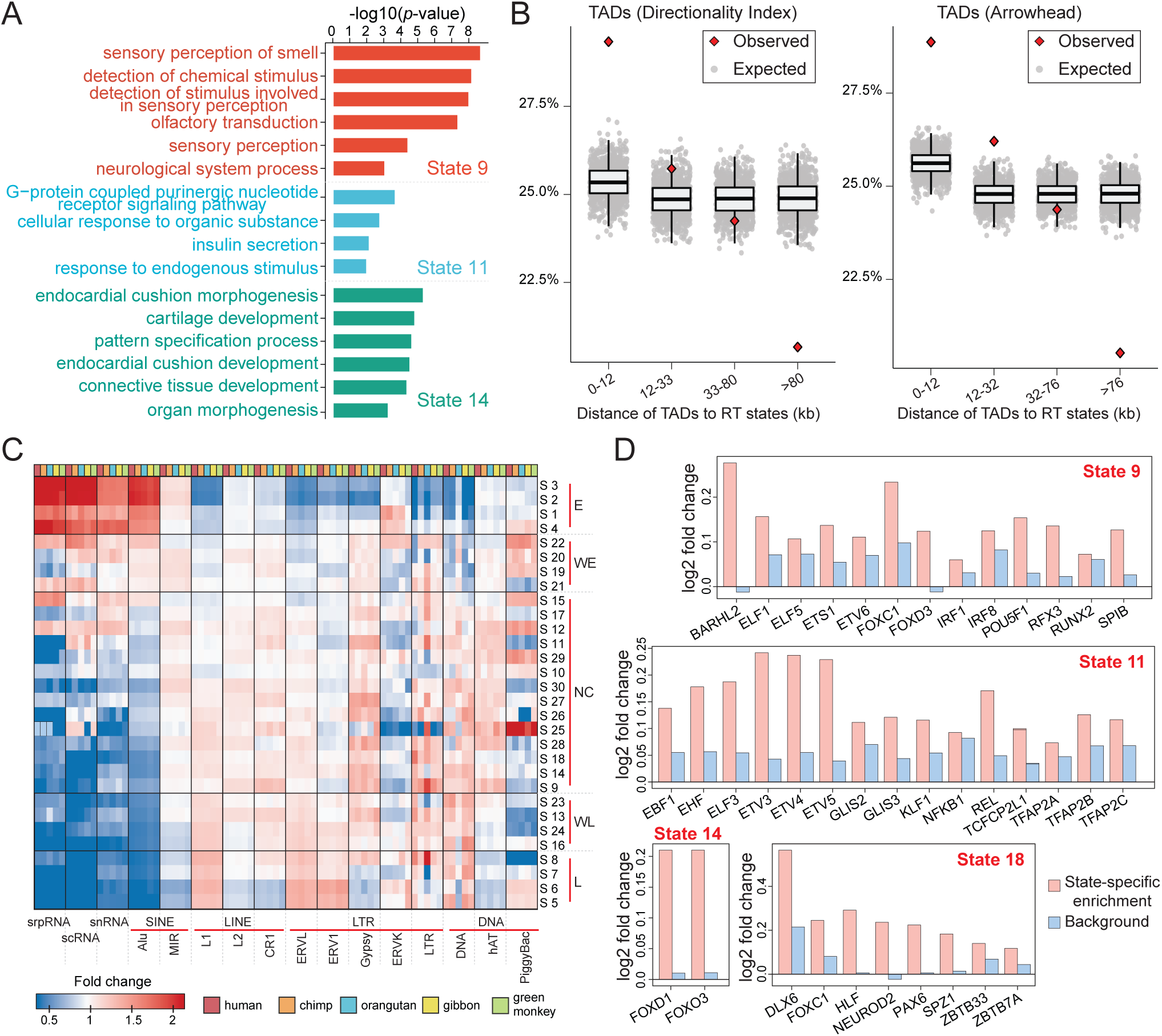
Comparisons between the RT evolution patterns and other genomic features. **(A)** Example gene ontology (GO) analysis results of state 9, state 11, and state 14. **(B)** Percentages of the distances between TAD boundaries and boundaries of predicted states in different intervals. The expected distances are calculated based on randomly shuffled TADs. Two types of TADs from different methods are used, namely TADs called by the Directionality Index method and TADs called by Arrowhead. **(C)** Transposable element enrichment in different RT states. **(D)** Motif enrichment in different lineage-specific RT states. State 9: human-chimpanzee specific early RT. State 11: human-chimpanzee-orangutan specific early RT. State 14: orangutan specific early RT. State 18: green monkey specific early RT. See also Fig. S2, Table S3, and Table S4. The GO analysis results of other lineage-specific RT states are included in Table S4.

### Boundaries of RT evolution patterns correlate with TAD boundaries

Earlier studies discovered that TADs defined from Hi-C data have high correspondence with replication domains (Pope et al., 2014; Dileep et al., 2015). We next asked whether the states found by Phylo-HMGP correlate with the boundaries of TADs. We used the TADs called by two methods, Directionality Index (DI) (Dixon et al., 2012) and Arrowhead (Rao et al., 2014). We named the TADs as DI TADs and Arrowhead TADs, the median lengths of which are 440kb and 185kb, respectively. For each boundary of a TAD, we calculated the distance between the TAD boundary and the nearest state boundary from Phylo-HMGP. We then calculated the percentages of boundary distances that fall into four distance intervals respectively, and estimated the empirical distributions of boundary distances in the different intervals by shuffling the TADs (Supplementary Methods A.10).

We found that the boundary distances between the DI TADs and the predicted RT states are significantly more enriched in the interval [0,12kb] than expected (Fig. 4B, empirical *p*-value<1e-03). The percentage drops in the intervals that correspond to increased boundary distances. The percentage is significantly lower than expected in the fourth interval that covers the largest distances (empirical *p*-value<1e-03). The comparison based on Arrowhead TADs show similar results. This analysis demonstrates the correlation between the boundaries of RT evolution states and the TAD boundaries.

### RT evolution patterns have enrichment of different transposable elements

It is known that RT correlates with certain transposable element (TE) families, e.g., the early RT regions are typically enriched with SINE elements (Rhind and Gilbert, 2013). We next looked at the connection between RT evolution patterns and the involvement of TEs based on RepeatMasker annotation. We obtained the RepeatMasker annotations for each of the five primate species from the UCSC genome browser (Karolchik et al., 2004). For the TE families shared among the five primate species, we calculated the fold change of their enrichment in the orthologous regions of each species in each state (Fig. 4C). We found that there exist distinct patterns of TE enrichment across different RT states and RT groups. Alu elements are strongly involved in conserved early RT states and depleted in conserved late RT states across the five species, with a clear changing correlation with RT across the five RT groups. On the contrary, L1 and LTR elements ERVL and ERV1 correlate negatively with early RT but positively with late RT. TEs in the LTR class and DNA class generally have more diversity in their distributions over states in the WE, WL, and NC groups. We also found that the repetitive sequence elements srpRNA, scRNA, and snRNA (based on RepeatMasker annotations) have a strong positive correlation with conserved early RT and negative correlation with conserved late RT (*p*-value<1e-04), having a similar enrichment pattern to Alu in the E, WL, and L groups. Although some of these correlations (such as those with srpRNA, scRNA, and snRNA) have not been reported before and further investigations are needed, this nevertheless demonstrates the potential of our method to provide new insights into the impact of sequence evolution on DNA replication timing.

### Lineage-specific early RT regions harbor unique TFBS

We then asked whether there are specific transcription factor binding sites (TFBS) that are enriched in regions with specific types of RT evolution patterns. We used FIMO (Grant et al., 2011) to perform motif scanning in the orthologous open chromatin regions of each species (Supplementary Methods A.11), using 635 position weight matrices (PWMs) of TF binding motifs from the JASPAR 2016 core vertebrate motif database (Mathelier et al., 2016). We then computed the motif frequency for each of the PWMs for each species, using the threshold of *p*-value<1e-04, and normalized the frequency by the open chromatin region size. We used two types of tests jointly to identify TF binding motifs that may be enriched in predicted lineage-specific RT states. First, within each lineage-specific RT state, we performed binomial tests to find the motifs that are more enriched in the particular lineage than expected (*p*-value<0.05). Second, we examined if the species-specific enrichment of a motif in a state is also significantly different from the genome-wide background distribution (Supplementary Methods A.11).

We identified a set of motifs that show lineage-specific enrichment for each of the lineage-specific early RT states (Fig. 4D). We found that the identified lineage-specific enriched TF motifs vary in different states. However, there are still a number of TF binding motifs (or motifs with similar PWMs) shared between different states. For example, FOXC1 is significantly enriched in human and chimpanzee in the hominini-specific state (state 9), and also enriched in green monkey in the green monkey-specific state (state 18). Interestingly, many of the TF motifs associated with species-specific early RT are from the FOX family (e.g., FOXC1, FOXO3, FOXD1, and FOXD3), the ELF family (e.g., ELF1 and ELF3), and the ETV family (e.g., ETV3 and ETV6). TFs of the FOX family are known regulators in B cells (Laurenti et al., 2013) (lymphoblastoid cells are B cells) and FOXO3 was previously found to be crucial for regulating cell cycle progression through its binding partnership with DNA replication factor Cdt1 (Zhang et al., 2012). Many of the other identified TFs are also regulators in B cells, such as EBF1, IRF8, RUNX2, and POU5F1 (Laurenti et al., 2013). Although these findings need further studies to evaluate their functional significance in lineage-specific biology, our analysis points to the direction that connects lineage-specific changes in *cis*-regulatory elements with lineage-specific changes in RT.

## Discussion

In this paper, we developed Phylo-HMGP, which is a new continuous-trait probabilistic model for more accurate genome-wide state estimation based on multivariate features from different species using functional genomic signals. The proposed Phylo-HMGP explores a new integrated framework to utilize the continuous-trait evolutionary model with spatial constraints to more effectively study the heterogeneous evolutionary feature patterns encoded in the genome-wide functional genomic datasets across multiple species. Both simulation studies and real data application demonstrate the advantage of Phylo-HMGP as compared to other methods. Importantly, we generated a new cross-species RT dataset from the same cell type in five primate species (human, chimpanzee, orangutan, gibbon, and green monkey) to study RT evolution patterns in primates for the first time using Phylo-HMGP. Our results from the comparative RT analysis demonstrate the potential of the model to help reveal regions with conserved or lineage-specific regulatory roles for the entire genome.

There are a number of areas that our model can be further improved. For Phylo-HMGP, the number of model parameters increases linearly with the number of species. There can be many local minima in parameter estimation for large scale evolutionary trees. Therefore, both more effective parameter constraints in accordance with the tree structure and more effective optimization methods need to be developed. Also, hierarchical state estimation methods can be developed to to group similar predicted patterns for state prediction refinement. In addition, the current Phylo-HMGP assumes that all the phylogenetic tree models have the same tree topolopy. But in certain application domains this may not be accurate. Therefore, it would be useful to improve the model by incorporating inference of alternative tree topologies (Friedman et al., 2002). Furthermore, we need to improve the interpretation of the estimated model parameters of the evolutionary models associated with the predicted states, to gain deeper understanding of the evolutionary mechanisms underlying the different functional genomic feature patterns.

We believe that Phylo-HMGP provides a generic framework to more precisely capture the evolutionary history of functional genomic signals across different species. In addition to the cross-species RT comparisons, we also applied Phylo-HMGP to predict the evolution of *cis-*regulatory modules and demonstrated the advantage and the generic utility of our new method (Supplementary Results B.2, Fig. S8, and Fig. S9). From the application to the RT data, we found that different RT evolution patterns predicted by Phylo-HMGP correlate with RT patterns across different cell types and various other genomic and epigenomic features, including higher-order genome organization features, *cis*-regulatory elements, transposons, and gene functions. Such insights from comparative functional genomic analysis may in turn help interpret the impact of sequence evolution on genome organization and function. One important future direction would be to develop more integrated models to holistically consider sequence features and functional genomic signals across multiple species.

## Methods

### Data simulation for the simulation studies

We used two types of models for data simulation, corresponding to Simulation Study I (SS-I) and Simulation Study II (SS-II) (Supplementary Methods A.5). Each study consists of six synthetic datasets. In SS-I, for each dataset, samples were generated from an HMM with 10 states and with multivariate Gaussian distribution as the emission probability distribution. The Gaussian distribution of each state follows a different OU model on the same phylogenetic tree topology. The OU model parameters of each state were randomly generated with non-negative constraints of selection strength 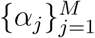 and Brownian motion intensity 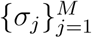 (*M* is the state number). Phylogenetic trees with four leaf nodes and with five leaf nodes were used as tree topologies for parameter simulation, each used for three datasets. The transition probability matrix of the HMM was randomly generated with the assumption that self-transition probability of a state is the dominant probability as compared to probabilities of transitions to other states.

In SS-II, samples were generated from a Gaussian mixture model instead of an HMM. For each dataset, 50,000 samples were generated based on a mixture model with 10 states where Gaussian distributions are the emission probability distributions. We defined a transition probability matrix between the 10 states as we defined in SS-I, and computed the equilibrium probability distribution of the 10 states from the transition probability matrix. We then divided the genome into continuous segments of varied lengths. Each segment represents a series of samples that share the same state, e.g., adjacent fixed-size bins of the same state on the genome. We randomly sampled the segment length from a truncated Normal distribution by which the length is non-negative. The state of each segment was drawn randomly and independently from the computed equilibrium probability distribution of the 10 states. Parameters of the Gaussian distribution of each state were shared between two corresponding datasets in SS-I and SS-II. For example, simulation dataset I-1 (dataset 1 in SS-I) and simulation dataset II-1 (dataset 1 in SS-II) are assigned with the same set of Gaussian distributions for 10 states. However, different assumptions (HMM and non-HMM models) were used to simulate the two types of datasets.

### Cell culture, replication timing profiling, and Repli-seq data processing

The GM12878 cells were grown according to the protocols defined in https://data.4dnucleome.org/biosources/4DNSRH17RFKR/ and https://data.4dnucleome.org/biosamples/4DNBS3I5U7BY/. For the other primate species, lymphoblastoid suspension cells were grown in RPMI-1640 media supplemented with 15% FBS and 2mM L-glutamine. Cultures were maintained in T25 flasks at 37°C, 5% CO_2_ and passaged to maintain adequate confluency. Next, we performed Repli-seq for the cultured cells of each species using the protocol defined in (Marchal et al., 2017).

To obtain multi-species RT signals from the Repli-seq data, we first mapped the preprocessed sequencing reads after quality control to the genome assemblies of hg19, panTro4, ponAbe2, nomLeu3, and chlSab1, respectively, using Bowtie2 (Langmead and Salzberg, 2012). Second, we used human genome (hg19) as the reference and divided the reference genome into 6kb bins. We then aligned each bin in human genome to each of the other species with reciprocal mapping using liftOver (Hinrichs et al., 2006) to obtain the orthologous regions. Third, for each species, we calculated Repli-seq read count within a given genomic window (an orthologous region) in early and late phases of RT, respectively, normalized by the total read count in early or late RT phase on the whole genome accordingly. The RT signal in each orthologous region is defined as the base 2 logarithm ratio of read count per million reads between the early and late phases of RT within this region (see Supplementary Methods A.7 for additional data processing steps). For each 6kb genomic bin in human, if the RT signals in the orthologous regions across five species are all available, we form the signals into a feature vector and assign it to the bin as a sample. Overall, we obtained 419,754 samples.

### Ornstein-Uhlenbeck process in the Phylo-HMGP model

#### Overall framework

We define a Phylo-HMGP model as **h** = (*S, ψ, A, π*), where *S* is the set of states, *ψ* is the set of phylogenetic models, *A* is the state-transition probability matrix, and *π* represents the initial state probabilities, respectively.

In Phylo-HMGP-OU, we can model the continuous traits with the Ornstein-Uhlenbeck process, which is a stochastic process that extends the Brownian motion with the trend towards equilibrium around optimal values. It is characterized by the following stochastic differential equation (Hansen, 1997; Butler and King, 2004):

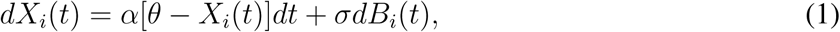

where *X*_*i*_(*t*) represents the observation of the *i*-th species at time point *t, B*_*i*_(*t*) is the Brownian motion, *α, θ*, and *σ* are parameters that represent the selection strength, the optimal value, and the Brownian motion intensity, respectively. For example, *X*_*i*_ could be the ChIP-seq signal of a certain histone mark from a specific cell type at a specific locus in a species. Under the assumption of the OU process, we can derive the expectation, the variance, and the covariance of the observations of species given the phylogenetic model *ψ*_*j*_. The phylogenetic model is the combination of multiple OU processes that share parameters along common branches. Suppose that *X*_*p*_ is the trait value of the ancestor of the *i*-th species, and *X*_*a*_ is the trait value of the common ancestor of the *i*-th and *j*-th species. Following Butler and King (2004); Rohlfs et al. (2013), we have:

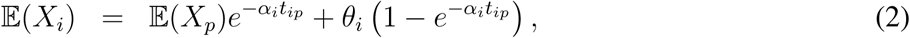

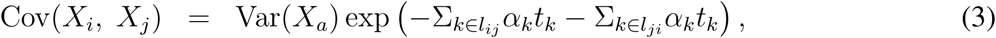

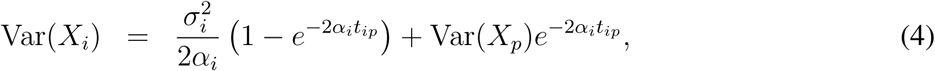

where *t*_*ip*_ is the length of the branch from *p* to *i*, and *l*_*ij*_ represents the set of the ancestor nodes of *i* and *i* itself after its divergence with *j*.

In the Phylo-HMGP model with OU process, *ψ*_*j*_ is defined as: *ψ*_*j*_ = (*θ*_*j*_, *α*_*j*_, *σ*_*j*_, *τ*_*j*_, *β*_*j*_), *j* = 1,…, *M*, where *θ*_*j*_, *α*_*j*_, *σ*_*j*_ denote the OU process parameters of the *j*-th state, respectively. *τ*_*j*_, *β*_*j*_ represent the topology of the phylogenetic tree and the branch lengths, respectively. We allow varied selection strength and Brownian motion intensity along each branch and varied optimal values at interior nodes or leaf nodes. Suppose there are *n* branches. We have *θ*_*j*_ ∈ ℝ^*n*+1^, *α*_*j*_, *σ*_*j*_ ∈ ℝ^*n*^. Suppose **x** = (*x*_1_, …, *x*_*N*_) are observations of consecutive regions along a sequence of length *N*, and **y** = (*y*_1_, …, *y*_*N*_) are the underlying hidden states, respectively. Each observation *x*_*i*_ is a multi-dimensional vector of the trait values of the compared species with respect to a certain type of functional genomic feature for an orthologous genomic region. Suppose there are *d* species, which correspond to the *d* leaf nodes in the phylogenetic tree. We have *x*_*i*_ ∈ ℝ^*d*^, *y*_*i*_ ∈ {1, …, *M*}, *i* = 1, …, *N*. The hidden state *y*_*i*_ indicates a specific phylogenetic model *ψ*_*j*_ from which the observation *x*_*i*_ is generated. 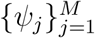 represent different evolutionary patterns of the genomic features across the multiple species. For example, one phylogenetic model *ψ*_*i*_ may represent conserved evolution of the feature across species, while another model *ψ*_*j*_ may represent strong selection strength along one lineage that results in a lineage-specific pattern. Given the input of multi-species functional genomic signals over a range of regions, which can be processed into the observations **x**, we are trying to infer the underlying evolutionary patterns and predict the evolutionary states **y** through model parameter estimation. Each *y*_*i*_ represents an evolutionary pattern, parameterized by the inferred phylogenetic model 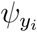. The output includes the estimated model parameters ĥ and predicted states **ŷ**. Note that our Phylo-HMGP-OU is different from the HMMSDE methods that use a temporal HMM to simulate a single OU process (Dittmer, 2009). Phylo-HMGP-OU embeds phylogenetic models constructed by complex of OU processes into a spatial HMM to utilize both temporal and spatial dependencies between variables.

#### Parameter estimation

Let Θ be the model parameters. The joint probability of the observations **x** and states **y** is 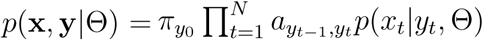 (Bilmes et al., 1998). We use Expectation-Maximization (EM) algorithm (Dempster et al., 1977) for parameter estimation. Suppose Θ^*g*^ is the current estimate of model parameters. The EM algorithm computes the expectation of the complete-data log likelihood, which is defined as the *Q* function *Q*(Θ, Θ^*g*^):

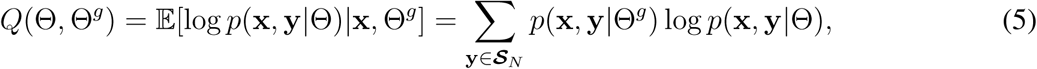

where ***𝒮***_*N*_ is the set of all state sequences of length *N*. We have:

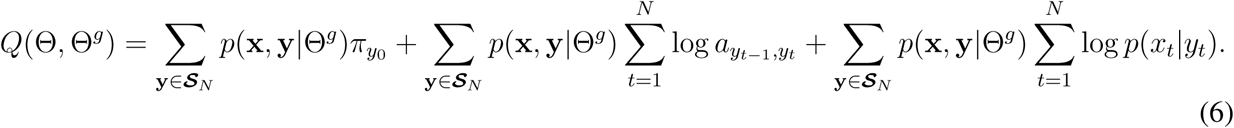

Model parameters *π, A* and can be updated separately in the Maximization-step (M-step), corresponding to the three parts of *Q*(Θ, Θ^*g*^), respectively. Define that *q*_*-t*_ = (*y*_1_,…, *y*_*t-*1_, *y*_*t*+1_, …, *y*_*N*_). We represent the third part as:

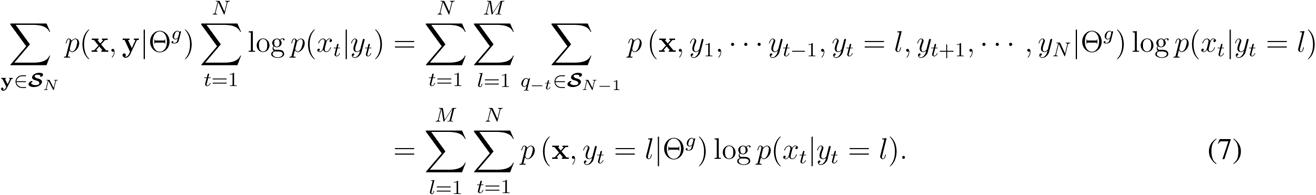

*p*(**x**, *y*_*t*_ = *l*|Θ^*g*^) can be computed using forward-backward algorithm (Rabiner, 1989; Bilmes et al., 1998). We have log 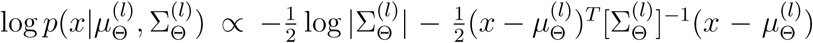 for a given state *l*. The underlying phylogenetic model *ψ*_*l*_ is embedded into 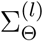 and 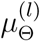 by Eq. (2)-(4). Assume that continuous-trait variables follow multi-variate Gaussian distributions. The negative log likelihood of state *l* is:

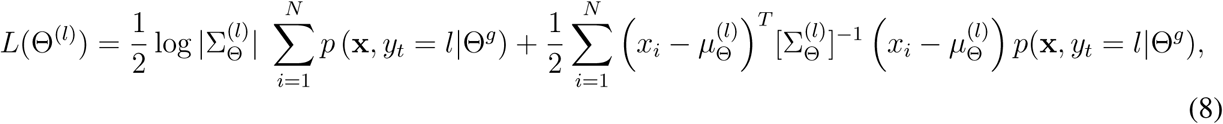

We need to perform parameter estimation for each of the possible states. We assume *τ*_*l*_ is given. *β*_*l*_ can be combined in effect to *α*_*l*_ and *σ*_*l*_. In practice, if the real branch lengths are unknown, we perform the transformation that 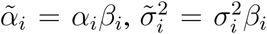, where *β*_*i*_ represents the length of branch from the parent of node *i* to node *i*. Using this approach the branch lengths are incorporated into {*α*_*l*_, *σ*_*l*_}. Then Θ^(*l*)^ = {*θ*_*l*_, *α*_*l*_, *σ*_*l*_}. A challenge is that there are approximately two times more model parameters than the feature dimension for each state. We apply ℓ_2_-norm regularization to the parameters Θ^(*l*)^. In each M-step, the objective function of a given state *l* is defined as:

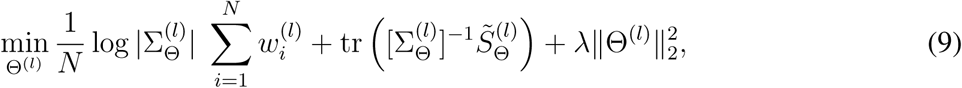

where 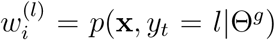 and 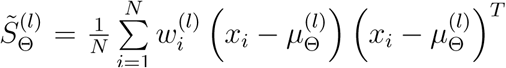. we define 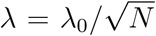, and tune *λ*_0_ based on a fixed simulation dataset. We estimated the range of *λ*_0_ that can improve the performance of Phylo-HMGP-OU (Supplementary Methods A.4 and Fig. S1). Accordingly, we applied the same *λ*_0_ to all the simulation datasets and the real data as a fixed coefficient, without tuning *λ*_0_ on each dataset specially, in order to avoid overfitting of *λ*_0_ on a particular dataset.

From the first two parts of *Q*(Θ, Θ^*g*^) we can update the estimates of ***π*** and *A* accordingly:

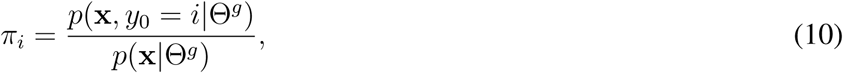

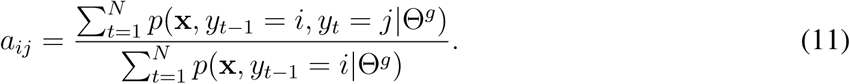

Therefore, in each E-step, given the estimated parameters Θ^*g*^, we compute *p*(**x**, *y*_*t*_ = *l*|Θ^*g*^) and *p*(**x**, *y*_*t-*1_ = *i, y*_*t*_ = *j*|Θ^*g*^) using the forward-backward algorithm, *l* = 1, …, *M*. In each M-step, we solve the Maximum Likelihood Estimation problem to update the parameters *π, A*, and 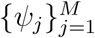. Given the estimated model parameters, we can predict a most likely sequence of hidden states **ŷ** using the Viterbi algorithm (Viterbi, 1967). Further details of Phylo-HMGP-OU are included in Supplementary Methods A.1 and A.3.

Note that the existing discrete-trait Phylo-HMMs (Siepel and Haussler, 2005; Hobolth et al., 2007) can also be represented as **h** = (*S, ψ, A, π*), where *ψ*_*j*_ is defined according to the substitution process with respect to an alphabet Σ_*j*_ of discrete characters, e.g., Σ_*j*_ = {*A, C, G, T*} for nucleotides. In discrete-trait Phylo-HMMs, *ψ*_*j*_ is defined as *ψ*_*j*_ = {*Q*_*j*_, *b*_*j*_, *τ*_*j*_, *β*_*j*_}, where *Q*_*j*_ is the substitution rate matrix, *b*_*j*_ is the vector of the background character frequencies, *τ*_*j*_ is the tree topology, and *β*_*j*_ represents the branch lengths. This realization of *ψ*_*j*_ is limited to the discrete characters, where transition probabilities between two characters can be computed to model evolution of characters, e.g., the HKY85 model (Hasegawa et al., 1985). For the continuous traits, we need to use continuous-trait evolutionary model assumptions to define *ψ*_*j*_.

### Brownian motion in the Phylo-HMGP model

For more comprehensive method evaluation of the proposed framework, we also developed the Phylo-HMGP-Brownian Motion (Phylo-HMGP-BM) method, where the embedded continuous-trait model is the Brownian motion model. Phylo-HMGP-BM is also built from **h** = (*S, ψ, A, π*). For Phylo-HMGP-BM, *ψ*_*j*_ is defined as *ψ*_*j*_ = (*µ*_*j*_, *τ*_*j*_, *β*_*j*_, *λ*_*j*_), 1 ≤*j* ≤ *M*, where *µ*_*j*_ denotes the mean values of leaf nodes, and *τ*_*j*_, *β*_*j*_, *λ*_*j*_ denote the phylogenetic tree topology, the branch lengths, and the evolution rates on branches, respectively. Under the Brownian motion assumption, the covariance between observations of two species depends on the depth of their nearest common ancestor in the phylogenetic tree. The covariance matrix based on the BM model can therefore be presented as a linear combination of covariance matrices (Zwiernik et al., 2017). More detailed description of Phylo-HMGP-BM is in Supplementary Methods A.2.

## Acknowledgements

This work has been supported primarily by the National Institutes of Health grant R01HG007352 (to J.M.). J.M. has been additionally supported by the National Science Foundation grants 1054309, 1262575, and 1717205. D.M.G. has been supported by the National Institutes of Health grant R01GM083337. J.M. and D.M.G. are also supported by the National Institutes of Health grant U54DK107965.

## Author Contributions

Conceptualization, J.M.; Methodology, Y.Y., Q.G., and J.M.; Software, Y.Y.; Investigation, Y.Y., Q.G., Y.Z., T.S., D.M.G., and J.M.; Resources, J.C. and R.J.O.; Visualization, Y.Z.; Writing – Original Draft, Y.Y. and J.M.; Writing – Review & Editing, Y.Y., Q.G., Y.Z., R.J.O., D.M.G., and J.M.; Funding Acquisition, D.M.G. and J.M.

## References

J. A. Bilmes et al. A gentle tutorial of the em algorithm and its application to parameter estimation for gaussian mixture and hidden markov models. 1998.

D. Brawand, M. Soumillon, A. Necsulea, P. julien, G. Csárdi, P. harrigan, M. Weier, A. Liechti, A. Aximu-Petri, M. Kircher, et al. The evolution of gene expression levels in mammalian organs. Nature, 478(7369):343–348, 2011.

M. A. Butler and A. A. King. Phylogenetic comparative analysis: a modeling approach for adaptive evolution. The American Naturalist, 164(6):683–695, 2004.

J. Cotney, J. Leng, J. Yin, S. K. Reilly, L. E. DeMare, D. Emera, A. E. Ayoub, P. rakic, and J. P. Noonan. The evolution of lineage-specific regulatory activities in the human embryonic limb. Cell, 154(1): 185–196, 2013.

A. P. dempster, N. M. Laird, and D. B. Rubin. Maximum likelihood from incomplete data via the em algorithm. Journal of the royal statistical society. Series B (methodological), pages 1–38, 1977.

V. Dileep, F. Ay, J. Sima, D. L. Vera, W. S. Noble, and D. M. Gilbert. Topologically associating domains and their long-range contacts are established during early g1 coincident with the establishment of the replication-timing program. Genome research, 2015.

E. Dittmer. Hidden Markov Models with time-continuous output behavior. PhD thesis, Freie Universität Berlin, 2009.

J. R. Dixon, S. Selvaraj, F. Yue, A. Kim, Y. Li, Y. Shen, M. Hu, J. S. Liu, and B. Ren. Topological domains in mammalian genomes identified by analysis of chromatin interactions. Nature, 485(7398): 376–380, 2012.

A. J. Drummond, M. A. Suchard, D. Xie, and A. Rambaut. Bayesian phylogenetics with beauti and the beast 1.7. Molecular biology and evolution, 29(8):1969–1973, 2012.

Q. Du, S. A. Bert, N. J. Armstrong, C. E. Caldon, J. Z. Song, S. S. Nair, C. M. Gould, P. L. Luu, A. Khoury, W. Qu, et al. Replication timing shapes the cancer epigenome and the nature of chromosomal rearrangements. bioRxiv, page 251280, 2018.

J. Felsenstein. Phylogenies and the comparative method. The American Naturalist, 125(1):1–15, 1985.

R. P. Freckleton. Fast likelihood calculations for comparative analyses. Methods in Ecology and Evolution, 3(5):940–947, 2012.

N. Friedman, M. Ninio, I. Pe’er, and T. Pupko. A structural em algorithm for phylogenetic inference. Journal of Computational Biology, 9(2):331–353, 2002.

C. E. Grant, T. L. Bailey, and W. S. Noble. Fimo: scanning for occurrences of a given motif. Bioinformatics, 27(7):1017–1018, 2011.

T. F. Hansen. Stabilizing selection and the comparative analysis of adaptation. Evolution, pages 1341–1351, 1997.

T. F. Hansen, J. Pienaar, and S. H. Orzack. A comparative method for studying adaptation to a randomly evolving environment. Evolution, 62(8):1965–1977, 2008.

M. Hasegawa, H. Kishino, and T.-a. Yano. Dating of the human-ape splitting by a molecular clock of mitochondrial dna. Journal of molecular evolution, 22(2):160–174, 1985.

A. S. Hinrichs, D. Karolchik, R. Baertsch, G. P. barber, G. Bejerano, H. Clawson, M. Diekhans, T. S. Furey, R. A. Harte, F. Hsu, et al. The ucsc genome browser database: update 2006. Nucleic acids research, 34(suppl 1):D590–D598, 2006.

A. Hobolth, O. F. Christensen, T. Mailund, and M. H. Schierup. Genomic relationships and speciation times of human, chimpanzee, and gorilla inferred from a coalescent hidden markov model. PLoS genetics, 3(2):e7, 2007.

D. W. Huang, B. T. Sherman, Q. Tan, J. R. Collins, W. G. Alvord, J. Roayaei, R. Stephens, M. W. Baseler, H. C. Lane, and R. A. Lempicki. The david gene functional classification tool: a novel biological module-centric algorithm to functionally analyze large gene lists. Genome biology, 8(9): R183, 2007.

J. L. Jensen and A.-M. K. Pedersen. Probabilistic models of dna sequence evolution with context dependent rates of substitution. Advances in Applied Probability, 32(2):499–517, 2000.

D. Karolchik, A. S. Hinrichs, T. S. Furey, K. M. Roskin, C. W. Sugnet, D. Haussler, and W. J. Kent. The ucsc table browser data retrieval tool. Nucleic acids research, 32(suppl 1):D493–D496, 2004.

B. Langmead and S. L. Salzberg. Fast gapped-read alignment with bowtie 2. Nature methods, 9(4):357, 2012.

E. Laurenti, S. Doulatov, S. Zandi, I. Plumb, J. Chen, C. April, J.-B. Fan, and J. E. Dick. The transcriptional architecture of early human hematopoiesis identifies multilevel control of lymphoid commitment. Nature immunology, 14(7):756, 2013.

M. Levin, L. Anavy, A. G. Cole, E. Winter, N. Mostov, S. Khair, N. Senderovich, E. Kovalev, D. H. Silver, M. Feder, et al. The mid-developmental transition and the evolution of animal body plans. Nature, 531(7596):637–641, 2016.

E. Lieberman-Aiden, N. L. Van Berkum, L. Williams, M. Imakaev, T. Ragoczy, A. Telling, I. Amit, B. R. Lajoie, P. J. Sabo, M. O. Dorschner, et al. Comprehensive mapping of long-range interactions reveals folding principles of the human genome. Science, 326(5950):289–293, 2009.

K. J. Liu, J. Dai, K. Truong, Y. Song, M. H. Kohn, and L. Nakhleh. An hmm-based comparative genomic framework for detecting introgression in eukaryotes. PLoS computational biology, 10(6):e1003649, 2014.

G. Lunter and J. Hein. A nucleotide substitution model with nearest-neighbour interactions. Bioinformatics, 20(suppl 1):i216–i223, 2004.

C. D. Manning, P. raghavan, and H. Schütze. Introduction to Information Retrieval. Cambridge University Press, New York, NY, USA, 2008. ISBN 0521865719, 9780521865715.

C. Marchal, T. Sasaki, D. Vera, K. Wilson, J. Sima, J.-C. Rivera-Mulia, C. T. Garcia, C. Nogues, E. Nafie, and D. M. Gilbert. Repli-seq: genome-wide analysis of replication timing by next-generation sequencing. bioRxiv, page 104653, 2017.

A. Mathelier, O. Fornes, D. J. Arenillas, C.-y. Chen, G. Denay, J. Lee, W. Shi, C. Shyr, G. Tan, R. Worsley-Hunt, et al. Jaspar 2016: a major expansion and update of the open-access database of transcription factor binding profiles. Nucleic acids research, 44(D1):D110–D115, 2016.

M. Naval-Saánchez, D. Potier, G. Hulselmans, V. Christiaens, and S. Aerts. Identification of lineage-pecific cis-regulatory modules associated with variation in transcription factor binding and chromatin activity using ornstein–uhlenbeck models. Molecular biology and evolution, 32(9):2441–2455, 2015.

M. Pagel. Inferring the historical patterns of biological evolution. Nature, 401(6756):877, 1999.

B. D. Pope, T. Ryba, V. Dileep, F. Yue, W. Wu, O. Denas, D. L. Vera, Y. Wang, R. S. Hansen, T. K. Canfield, et al. Topologically associating domains are stable units of replication-timing regulation. Nature, 515(7527):402–405, 2014.

J. Qu, E. Hodges, A. Molaro, P. gagneux, M. D. Dean, G. J. Hannon, and A. D. Smith. Evolutionary expansion of dna hypomethylation in the mammalian germline genome. Genome research, 28(2): 145–158, 2018.

L. R. Rabiner. A tutorial on hidden markov models and selected applications in speech recognition. Proceedings of the IEEE, 77(2):257–286, 1989.

S. S. Rao, M. H. Huntley, N. C. Durand, E. K. Stamenova, I. D. Bochkov, J. T. Robinson, A. L. Sanborn, I. Machol, A. D. Omer, E. S. Lander, et al. A 3d map of the human genome at kilobase resolution reveals principles of chromatin looping. Cell, 159(7):1665–1680, 2014.

N. Rhind and D. M. Gilbert. Dna replication timing. Cold Spring Harbor perspectives in biology, 5(8):a010132, 2013.

J. C. Rivera-Mulia, Q. Buckley, T. Sasaki, J. Zimmerman, R. A. Didier, K. Nazor, J. F. Loring, Z. Lian, S. Weissman, A. J. Robins, et al. Dynamic changes in replication timing and gene expression during lineage specification of human pluripotent stem cells. Genome research, 25(8):1091–1103, 2015.

R. V. Rohlfs, P. harrigan, and R. Nielsen. Modeling gene expression evolution with an extended ornstein–uhlenbeck process accounting for within-species variation. Molecular biology and evolution, 31(1):201–211, 2013.

T. Ryba, I. Hiratani, J. Lu, M. Itoh, M. Kulik, J. Zhang, T. C. Schulz, A. J. Robins, S. Dalton, and D. M. Gilbert. Evolutionarily conserved replication timing profiles predict long-range chromatin interactions and distinguish closely related cell types. Genome Research, 20(6):761–770, 2010.

T. Ryba, I. Hiratani, T. Sasaki, D. Battaglia, M. Kulik, J. Zhang, S. Dalton, and D. M. Gilbert. Replication timing: a fingerprint for cell identity and pluripotency. PLoS computational biology, 7(10):e1002225, 2011.

T. Ryba, D. Battaglia, B. H. Chang, J. W. Shirley, Q. Buckley, B. D. Pope, M. Devidas, B. J. Druker, and D. M. Gilbert. Abnormal developmental control of replication-timing domains in pediatric acute lymphoblastic leukemia. Genome research, 22(10):1833–1844, 2012.

A. Siepel and D. Haussler. Phylogenetic hidden markov models. In Statistical methods in molecular evolution, pages 325–351. Springer, 2005.

I. Solovei, K. Thanisch, and Y. Feodorova. How to rule the nucleus: divide et impera. Current opinion in cell biology, 40:47–59, 2016.

D. Villar, C. Berthelot, S. Aldridge, T. F. Rayner, M. Lukk, M. Pignatelli, T. J. Park, R. Deaville, J. T. Erichsen, A. J. Jasinska, et al. Enhancer evolution across 20 mammalian species. Cell, 160(3):554–566, 2015.

N. X. Vinh, J. Epps, and J. Bailey. Information theoretic measures for clusterings comparison: Variants, properties, normalization and correction for chance. Journal of Machine Learning Research, 11(Oct): 2837–2854, 2010.

A. Viterbi. Error bounds for convolutional codes and an asymptotically optimum decoding algorithm. IEEE transactions on Information Theory, 13(2):260–269, 1967.

E. Yaffe, S. Farkash-Amar, A. Polten, Z. Yakhini, A. Tanay, and I. Simon. Comparative analysis of dna replication timing reveals conserved large-scale chromosomal architecture. PLoS genetics, 6(7): e1001011, 2010.

F. Yue, Y. Cheng, A. Breschi, J. Vierstra, W. Wu, T. Ryba, R. Sandstrom, Z. Ma, C. Davis, B. D. Pope, et al. A comparative encyclopedia of dna elements in the mouse genome. Nature, 515(7527):355, 2014.

Y. Zhang, Y. Xing, L. Zhang, Y. Mei, K. Yamamoto, T. W. Mak, and H. You. Regulation of cell cycle progression by forkhead transcription factor foxo3 through its binding partner dna replication factor cdt1. Proceedings of the National Academy of Sciences, 109(15):5717–5722, 2012.

P. Zwiernik, C. Uhler, and D. Richards. Maximum likelihood estimation for linear gaussian covariance models. Journal of the Royal Statistical Society: Series B (Statistical Methodology), 79(4):1269–1292, 2017.

