## Supplementary Materials for "Continuous-trait probabilistic model for comparing multi-species functional genomic data"

### Supplemental Information

#### A Supplementary Methods

##### A.1 Phylo-HMGP model with Ornstein-Uhlenbeck process

We use EM algorithm for parameter estimation in Phylo-HMGP-OU. The Q function $Q(\Theta, \Theta^g)$ is defined in Eq. (6). From the first two terms of $Q(\Theta, \Theta^g)$ we can update the estimates of $\boldsymbol{\pi}$ and $A$, as shown in Eq. (10) and Eq. (11).

The parameters of the Ornstein-Uhlenbeck (OU) model are involved in $p(x_t|y_t)$ of the third term of $Q(\Theta, \Theta^g)$. We have

$$\begin{aligned}
\sum_{\mathbf{y}\in\boldsymbol{\mathcal{S}}_N} p(\mathbf{x},\mathbf{y}|\Theta^g)\sum_{t=1}^{N}\log p(x_t|y_t) &= \sum_{t=1}^{N}\sum_{\mathbf{y}\in\boldsymbol{\mathcal{S}}_N} p(\mathbf{x},\mathbf{y}|\Theta^g)\log p(x_t|y_t)\\
&= \sum_{t=1}^{N}\sum_{l=1}^{M}\sum_{q_{-t}\in\boldsymbol{\mathcal{S}}_{N-1}} p(\mathbf{x},y_1,\cdots y_{t-1},y_t=l,y_{t+1},\cdots,y_N|\Theta^g)\log p(x_t|y_t=l)\\
&= \sum_{l=1}^{M}\sum_{t=1}^{N} p(\mathbf{x},y_t=l|\Theta^g)\log p(x_t|y_t=l) \qquad (12)
\end{aligned}$$

The third part of the negative log likelihood function with respect to a given state $l$ is:

$$\tilde{L}(\Theta^{(l)}) = \log|\Sigma_\Theta^{(l)}|\sum_{i=1}^{N} w_i^{(l)} + \operatorname{tr}\left([\Sigma_\Theta^{(l)}]^{-1}\tilde{S}_\Theta^{(l)}\right), \qquad (13)$$

where $w_i^{(l)} = p(\mathbf{x}, y_t = l|\Theta^g)$ and $\tilde{S}_\Theta^{(l)} = \frac{1}{N}\sum_{i=1}^{N} w_i^{(l)}\left(x_i - \mu_\Theta^{(l)}\right)\left(x_i - \mu_\Theta^{(l)}\right)^T$.

We apply $l_2$-norm regularization to the parameters $\Theta$. The objective function to maximize in the M-step of the EM algorithm is defined in Formula (9).

We compute $w_i^{(l)} = p(\mathbf{x}, y_t = l|\Theta^g)$ in the E-step using the forward-backward algorithm (Rabiner, 1989; Bilmes et al., 1998) and current estimates of the model parameters $\Theta_g$. We define:

$$\alpha_l(t) = p(x_1, x_2, \cdots, x_t, y_t = l|\Theta^g), \qquad (14)$$

and

$$\beta_l(t) = p(x_{t+1}, x_{t+2}, \cdots, x_T|y_t = l, \Theta^g). \qquad (15)$$

According to the forward-backward algorithm, we have:

$$p(\mathbf{x}, y_t = l|\Theta^g) = \alpha_l(t)\beta_l(t). \qquad (16)$$

Both $\alpha_j(t)$ and $\beta_j(t)$ can be computed recursively. Let $\pi_l = p(y_1 = l)$ be the initial state distribution, $l = 1, \cdots, M$. Let $A = \{a_{ij}\}$ be the transition matrix, where $a_{ij} = p(y_t = j|y_{t-1} = i)$. Let $X =$

$(x_1, \cdots, x_T)$ be the observation sequence. The forward procedure to compute $\alpha_l(t)$ is as follows.

$$\alpha_l(1) = \pi_l p(x_1|y_1 = l), \tag{17}$$

$$\alpha_l(t+1) = [\sum_{j=1}^{M} \alpha_j(t) a_{lj}] p(x_{t+1}|y_{t+1} = l), \tag{18}$$

$$p(X|\Theta) = \sum_{l=1}^{M} \alpha_l(T). \tag{19}$$

The backward procedure to compute $\beta_l(t)$ is as follows:

$$\beta_l(T) = 1, \tag{20}$$

$$\beta_l(t) = \sum_{j=1}^{M} a_{lj} y(x_{t+1}|y_{t+1} = j) \beta_j(t+1), \tag{21}$$

$$p(X|\Theta) = \sum_{l=1}^{M} \beta_l(1) \pi_l p(x_1|y_1 = l). \tag{22}$$

We also update the transition probability between any two states. Define that $\epsilon_{ij}(t) = p(y_t = i, y_{t+1} = j|X, \Theta)$. We have:

$$\epsilon_{ij}(t) = \frac{p(y_t = i, y_{t+1} = j, X|\Theta)}{p(X|\Theta)} = \frac{\alpha_i(t) a_{ij} p(x_{t+1}|y_{t+1} = j) \beta_j(t+1)}{\sum_{i=1}^{N} \sum_{j=1}^{N} \alpha_i(t) a_{ij} p(x_{t+1}|y_{t+1} = j) \beta_j(t+1)}. \tag{23}$$

The transition matrix can be updated as:

$$a_{ij} = \frac{\sum_{t=1}^{T-1} \epsilon_{ij}(t)}{\sum_{t=1}^{T-1} p(y_t = i|X, \Theta)}, i, j = 1, \cdots, M, i \neq j. \tag{24}$$

Therefore, in each E-step, given the estimated parameters $\Theta^g$, we compute $p(\mathbf{x}, y_t = l|\Theta^g)$ using the forward-backward algorithm, $l = 1, \cdots, M$. In each M-step, we solve the Maximum Likelihood Estimation problem to update the parameters associated with each state. Different optimization algorithms can be used to solve the optimization problem.

#### A.2 Phylo-HMGP model with Brownian motion

For Phylo-HMGP-BM, $\psi_j$ is defined as $\psi_j = (\mu_j, \tau_j, \beta_j, \lambda_j)$, $1 \leq j \leq M$, where $\mu_j$ denotes the mean values of leaf nodes, and $\tau_j$, $\beta_j$, $\lambda_j$ denote the phylogenetic tree topology, the branch lengths, and the evolution rate on each branch, respectively. Under the assumption of Brownian motion, the covariance between observations of two species is proportional to the depth of their nearest common ancestor in the phylogenetic tree. Suppose $X_i$ is the observation of a species. Based on the model of Brownian motion, the mean value of $X_i$ is identical to that of the observation of its ancestor and the variance of $X_i$ is proportional to the evolution time from its ancestor. Let $v_j = \lambda_j \beta_j$, which reflects the combined effect of branch length and evolution rate along this branch. We have:

$$\mathbb{E}[X_i] = \mathbb{E}[X_p], \tag{25}$$

$$\text{var}(X_i) = \sum_{k \in S_a(i)} v_k, \tag{26}$$

$$\text{cov}(X_i, X_j) = \sum_{k \in S_a(i,j)} v_k, \tag{27}$$

where $S_a(i)$ represents the set of ancestors of species $i$, and $S_a(i,j)$ represents the set of common of ancestors of species $i$ and $j$. The covariance matrix based on the Brownian motion model can therefore be presented as (Zwiernik et al., 2017):

$$\Sigma_v = G_0 + \sum_{i=1}^{r} v_i G_i, \tag{28}$$

where $G_i$ is a binary matrix representing contribution of a specific branch to the covariance matrix. Similar to Phylo-HMGP-OU, we use EM algorithm for parameter estimation. We define the $Q$ function in the same way as Eq. (6), which is the expectation of the complete-data log likelihood function, but with different realization of $p(\mathbf{x}, \mathbf{y}|\Theta^g)$ according to the assumption of the Brownian Motion model. Based on the original Brownian motion model, the expectation of observation of descendant species is always identical to that of its ancestor. We have $\mathbb{E}(X_i) = \mathbb{E}(X_0)$ under this assumption, where $X_0$ corresponds to the most remote ancestor. However, we can observe shift of the mean value of the phenotype in real world problems (Thomas et al., 2006, 2009). Therefore we relax this constraint on the expectation of the observations, using a weaker assumption that allows the expectation to be shifted on branches. Then we considers the phenotype expectation of each species as model parameters, allowing the expectation to vary between species.

Suppose $\Theta^g$ is the current estimate of the parameters. The objective function of a given state $l$ is:

$$\min_{v^{(l)}, \mu^{(l)}} \frac{1}{N} \log |\Sigma_{v,\mu}^{(l)}| \sum_{i=1}^{N} w_i^{(l)} + \operatorname{tr}\big([\Sigma_{v,\mu}^{(l)}]^{-1} \tilde{S}_\mu^{(l)}\big), \tag{29}$$

where $w_i^{(l)} = p(\mathbf{x}, y_t = l|\Theta^g)$, and $\tilde{S}_\mu^{(l)} = \frac{1}{N}\sum_{i=1}^{N} w_i^{(l)} \left(x_i - \mu^{(l)}\right)\left(x_i - \mu^{(l)}\right)^T$. Using EM algorithm, in each E-step, given the estimated parameters $\Theta^g$, we compute $p(\mathbf{x}, y_t = l|\Theta^g)$ using the forward-backward algorithm, $l = 1, \cdots, M$. In each M-step, we solve the Maximum Likelihood Estimation problem to update the parameters associated with each state. Different optimization algorithms can be applied to solve the optimization problem. For the Phylo-HMGP-BM, the gradient with respect to $v_j$ can be computed explicitly (Zwiernik et al., 2017) and we implemented the gradient descent method based on the derived gradient as an alternative optimization approach:

$$\frac{\partial \tilde{L}^{(l)}(v)}{\partial v_j^{(l)}} = \nabla_{G_j} \tilde{L}^{(l)}(v) = \frac{1}{2}\Sigma_{i=1}^{N} w_i^{(l)} \operatorname{tr}(G_j[\Sigma_{v,\mu}^{(l)}]^{-1}) + \frac{N}{2}\operatorname{tr}(\tilde{S}_N[\Sigma_{v,\mu}^{(l)}]^{-1} G_j [\Sigma_{v,\mu}^{(l)}]^{-1}). \tag{30}$$

##### A.3 Initialization of the Expectation-Maximization (EM) algorithm in Phylo-HMGP

Phylo-HMGP uses the EM algorithm for parameter estimation. The EM algorithm seeks local minima and the results of the EM algorithm are influenced by initializations. We designed different ways for parameter initialization. The first approach is to estimate OU model parameters initially based on the primitive state estimation results from K-means clustering. We perform model estimation for each cluster separately as single-state estimation, and use the estimates as initial model parameter values for the EM algorithm.

The second approach is to generate initial values randomly. There are three types of parameters in the OU model for a single state, which are optimal values $\theta$, Brownian motion intensity $\sigma$ and selection strength $\alpha$. We sample random variables from uniform distributions for the initial values of $\theta$, $\sigma$, and $\alpha$, respectively.

The third approach is to use linear combination of the initial parameter values obtained from the first approach and the second approach. We estimate the initial parameter values as $\Theta_0 = w_1\Theta_1 + (1 - w_2)\Theta_2$, where $\Theta_1$ and $\Theta_2$ are parameter estimates from the first and second approaches. By changing the initial weight $w_1$, we have different initialization schemes. Based on the performance with respect to varied $w_1$ in simulation study I, we observed that Phylo-HMGP is not very sensitive to initialization on four datasets, while on the other datasets the performance is improved as $w_1$ increases within a range. Given $w_1 \in [0.2, 1.0]$, the performance of Phylo-HMGP on each simulated dataset is comparable to the best performance it can achieve on the corresponding dataset. For performance comparison with other methods in the simulation study, we fixed $w_1 = 0.8$ for all the datasets to prevent overfitting on a particular dataset. The initialization weight $w_1$ is an input parameter to the implemented program and can be adjusted within $[0, 1]$ by the user's choice.

##### A.4 Estimation of the regularization coefficient in Phylo-HMGP

For the objective function defined in Formula (9), we define $\lambda = \lambda_0/\sqrt{N}$, where $N$ is the sample size, and we observe how performance of Phylo-HMGP-OU changes with respect to $\lambda_0$ based on a fixed simulation dataset (simulation dataset I-1), in order to estimate a range of $\lambda_0$ in which the performance of Phylo-HMGP-OU can be improved with the $l_2$-norm regularization. We tuned $\lambda_0$ from 0 to 5, with the step size of 0.5, and compared the performance of the model with respect to the different choices of $\lambda_0$. We found that the model with $\lambda_0 \in [3.0, 5.0]$ reaches relatively higher $F_1$ score than the other choices of $\lambda_0$ on this dataset (Fig. S1). We selected $\lambda_0 = 4.0$ and applied it to all the simulation datasets and the RT data as a fixed coefficient, without tuning $\lambda_0$ on each dataset specially, in order to avoid overfitting of $\lambda_0$ on a particular dataset. We also repeated the experiment on dataset I-1 and observed how the performance of Phylo-HMGP-OU changes with $\lambda_0$ on the other datasets in simulation study I. We found that the performance of Phylo-HMGP-OU is not sensitive to $\lambda_0$ ranging in [3.0,5.0] on most of the simulation datasets (I-1,I-3,I-4,I-5,I-6). Phylo-HMGP-OU still reaches comparable performance to the highest performance it can achieve on dataset II-2. We only used the performance resulted from $\lambda_0 = 4.0$ on all the simulation datasets for performance evaluation and comparison.

##### A.5 Data simulation for the simulation studies

In both simulation study I (SS-I) and II (SS-II), phylogenetic trees with four leaf nodes and with five leaf nodes were used as tree topologies for parameter simulation, each used for three datasets in each study. Datasets with even-number (I-2, I-4, I-6, II-2, II-4, and II-6) were based on the same topology of five leaf nodes, which is identical to the topology of the species tree specified in Fig. 1C and is also the topology used in the RT data study. Datasets with odd-number (I-1, I-3, I-5, II-1, II-3, and II-5) were based on the same topology of four leaf nodes, which is identical to the topology of the sub-tree of the species tree specified in Fig. 1C that contains human, chimpanzee, orangutan and gibbon.

The emission probability distribution of each state in each dataset is Gaussian distribution, parameterized by a multivariate OU model $\psi_j = (\theta_j, \alpha_j, \sigma_j)$, where $j$ is the index of the state, and $\theta_j$, $\alpha_j$, $\sigma_j$ represent the optimal value vector, the selection strengths and the Brownian motion intensities along the branches, respectively. The selection strength $\alpha_{j,k}$ and Brownian motion intensity $\sigma^2_{j,k}$ along each branch are each randomly and independently sampled from the uniform distribution $Unif[0, 2]$, $k = 1, \cdots, d$ ($d$ is the number of branches). The optimal value $\theta_{j,l}$ ($l = 1, \cdots, d + 1$) of each node is randomly and independently sampled from a Normal distribution $\mathcal{N}(0, 2)$.

In the transition probability matrix $A$ of one dataset in SS-I, the self-transition probability of state $j$ is defined as $a_{jj} = a_0 + (1 - a_0) \times p$, where $p$ is randomly sampled from uniform distribution $Unif[0, 1]$ and $a_0$ is set to be 0.7. The transition probabilities of state $j$ to other states are first randomly sampled from uniform distribution $Unif[0, 1]$ and then normalized to be summed to $1 - a_{jj}$.

In SS-II, the fragment length (the number of continuous bins of the same state) is sampled from the a truncated Normal distribution $\mathcal{N}(50, 30)$ with the minimal fragment length to be 5. We first sampled a transition probability matrix $\tilde{A}$ in the same way as in SS-I. Then we estimated the equilibrium probability distribution $\tilde{\pi}$ of the states from $\tilde{A}$ based on $\tilde{\pi} = \tilde{\pi}\tilde{A}$. We sampled the state of each fragment from $\tilde{\pi}$ randomly and independently.

We calculated the Davies-Bouldin Index (DBI) (Davies and Bouldin, 1979) for each dataset in SS-I and SS-II, to estimate the difficulty in state prediction in different datasets. DBI can be used to measure how discriminative is each cluster (state) compared to the others. A high DBI represents that the states have large variances within themselves while the state-to-state distances are small, making it difficult to distinguish the states. The DBIs for the six datasets in SS-I are 2.3127, 2.0770, 1.9706, 1.3127, 1.5045 and 1.4623, respectively. The DBIs for datasets in SS-II are 2.2677, 2.0608, 1.9597, 1.3116, 1.4987, and 1.4864, respectively. We found that datasets I-1, I-2, II-1, and II-2 have relatively higher DBIs.

##### A.6 Performance evaluation in the simulation studies

We used Adjusted Mutual Information (AMI), Normalized Mutual Information (NMI), Adjusted Rand Index (ARI), Precision, Recall and $F_1$ score (Manning et al., 2008; Vinh et al., 2010) for performance evaluation in the simulation studies. Suppose $X = \{x_1, \cdots, x_N\}$ is the set of samples. Suppose $\Omega = \{\omega_1, \cdots, \omega_K\}$ is the set of predicted states which represents a partition of $S$ into $K$ states, and $C = \{c_1, \cdots, c_M\}$ is the ground truth set of states. Let $I(\Omega, C)$ be the mutual information between $\Omega$ and $C$, and $NMI(\Omega, C)$ be the normalized mutual information. We have:

$$I(\Omega; C) = \sum_{k=1}^{K} \sum_{j=1}^{M} P(\omega_k, c_j) \log \frac{P(\omega_k, c_j)}{P(\omega_k)P(c_j)}, \tag{31}$$

$$NMI(\Omega; C) = \frac{I(\Omega; C)}{[H(\Omega) + H(C)]/2}, \tag{32}$$

where $H(\Omega)$ and $H(C)$ represent the entropies of $\Omega$ and $C$, respectively. The entropy is defined as $H(\Omega) = -\sum_{k=1}^{K} P(\omega_k) \log P(\omega_k)$. $P(\omega_k)$, $P(c_j)$, and $P(\omega_k, c_j)$ represent the probabilities that a sample is in state $\omega_k$, in state $c_j$, and in both $\omega_k$ and $c_j$, respectively. The maximum likelihood estimates of $P(\omega_k)$, $P(c_j)$, and $P(\omega_k, c_j)$ are $|\omega_k|/N$, $|c_j|/N$, and $|\omega_k \cap c_j|/N$, respectively, where $|\omega_k|$ denotes the size of $\omega_k$ and $N$ is the number of samples.

Adjusted Mutual Information (AMI) is an adjustment of the mutual information to correct the effect of agreement between two partitions that is solely due to chance. We have:

$$AMI(\Omega; C) = \frac{I(\Omega; C) - \mathbb{E}[I(\Omega; C)]}{\max\{H(\Omega), H(C)\} - \mathbb{E}[I(\Omega; C)]}, \tag{33}$$

where $\mathbb{E}(I(\Omega; C))$ represents the expectation of $I(\Omega; C)$, which can be estimated using $\Omega$ and $C$ (Vinh et al., 2010).

The Rand Index (RI) (Manning et al., 2008) is another metric to compare two partitions, which is defined as:

$$RI = \frac{TP + TN}{TP + FP + FN + TN}. \tag{34}$$

TP (true positive) represents the number of pairs of samples in $X$ that are in the same subset in $\Omega$ and also in the same subset in $C$. FP (false positive) is the number of pairs of samples in $X$ that are in the same subset in $\Omega$ but in different subsets in $C$. FN (false negative) is the number of pairs of samples in

$X$ that are in different subsets in $\Omega$ but in the same subset in $C$. TN (true negative) is the number of pairs of samples in $X$ that are in different subsets in $\Omega$ and also in different subsets in $C$.

The Adjusted Rand Index (ARI) corrects the Rand Index for the effect of agreement that is solely due to chance between partitions. ARI is defined as

$$ARI = \frac{RI - \mathbb{E}[RI]}{\max\{RI\} - \mathbb{E}[RI]}, \tag{35}$$

where $\mathbb{E}(RI)$ represents the expectation of $RI$.

Precision, Recall, and $F_1$ score are defined as

$$Precision = \frac{TP}{TP + FP}, \tag{36}$$

$$Recall = \frac{TP}{TP + FN}, \tag{37}$$

$$F_1 = \frac{2Precision \times Recall}{Precision + Recall}. \tag{38}$$

For the compared methods, we used the functions GaussianMixture and KMeans in the scikit-learn library (Pedregosa et al., 2011) to implement the Gaussian Mixture Model (GMM) method and the $K$-means clustering method, respectively. We used the hmmlearn library (https://github.com/hmmlearn/) to implement the Gaussian-HMM method. Each compared method is repeated 10 times with different initializations and guaranteed convergence each time. Specifically, the parameter initializations of the Gaussian-HMM method and the GMM method were based on $K$-means clustering. In each experiment, the best result from 10 randomly-initialized $K$-means clustering results (based on the clustering evaluation criteria used in hmmlearn or scikit-learn, respectively) was selected for estimating the initial parameters of Gaussian-HMM or GMM, respectively. 10 random initializations were also used for $K$-means clustering in each experiment, and the clustering with the best performance was chosen as the result. 10 experiments were repeated for the compared methods as well as Phylo-HMGP, and the average of the 10 runs was reported as the final performance of the corresponding method.

#### A.7 Data processing of the Repli-seq data

To process the raw Repli-seq data and obtain RT signals in orthologous genome regions across the multiple species, we computed the RT signal values for each 6kb bin of human genome and its orthologous regions in each of the other species if RT measurements are available. For data preprocessing, we performed quality control of the Repli-seq reads and removed adapter sequences using FASTX-Toolkit (http://hannonlab.cshl.edu/fastx_toolkit/). We used human genome (hg19) as the reference and divided the reference genome into 6kb bins, and performed alignment and RT signal calculation as described in the Methods section.

For each species, we identify each sequence of consecutive bins without RT signals as a gap. The bin size (6kb in human) is much smaller than the scale of the RT signals (the replication domain is typically at the scale of 400-800kb (Pope et al., 2014)). We assume that RT does not change sharply at a small size gap if the gap is between both early RT signals or both late RT signals. We then performed data imputation for gaps smaller than 48kb using nearest neighbor imputation. In this way we can reduce missing data and have more continuous segments where cross-species observations are available. More specifically, if the RT signals on both sides of a gap smaller than 48kb are both early RT signals or both late RT signals with difference smaller than 1/3, we assign to each bin in the gap the RT signal of a signal-available bin that is nearest to this bin. We then used the software HMMSeg (Day et al., 2007) to perform wavelet smoothing (Percival and Walden, 2006) of the RT signals in each species, using the window size of 24kb.

Next, we found the orthologous regions where the RT signals across five species are all available. We then performed data normalization of the signals of each species in the regions. We observed that the different species have varied RT scales around [-5,5]. We performed feature scaling to scale non-negative RT signals (primarily early RT) in each species to [0,5] and scale non-positive RT signals (primarily late RT) in each species to [-5,0]. We formed the normalized RT signals in orthologous regions across five species into a five-dimensional feature vector and assigned it to the corresponding reference 6kb bin in human genome as a sample. We obtained 419,754 samples in the orthogolous regions across species.

##### A.8 Initial estimation of the state number in the RT data study

To apply Phylo-HMGP to the replication timing data, we first estimated the possible number of states using K-means clustering. We performed K-means clustering to the datasets with an increasing cluster number $K$, computed the Sum of Squared Error (SSE) of each clustering result, and observed how SSE changed with respect to $K$. We estimated the state number to be approximately 20-40 based on the K-means clustering results, as the decreasing rate of SSE with respect to the increasing $K$ slows down in this range (Fig. S3). Based on the observation the 'elbow point' of the SSE curve is around 30, and small fluctuation of the state number around 30 dose not present significant change of the reduction of the SSE decreasing rate compared to the state number of 30. We therefore set the state number to be 30.

##### A.9 RT state prediction and RT state grouping

We applied Phylo-HMGP-OU to the multi-species Repli-seq data to perform state estimation, with the state number set to be 30. We used $\lambda_0 = 4.0$ for $l_2$-norm regularization and $w_1 = 0.2$ for parameter initialization (see Supplementary Methods A.3). We repeated the estimation 10 times with different initializations. We choose the result with the highest objective function value for further analysis.

We classified the 30 RT states predicted by Phylo-HMM-OU into 5 RT groups, namely, conserved early (noted as E), conserved late (L), weakly conserved early (WE), weakly conserved late (WL), and non-conserved (NC). If the majority (>98%) of the regions in a state share the pattern that all of the five species consistently have positive RT signals (early in RT), we assign this state to the conserved early (E) group. If a state does not satisfy this criteria, but instead satisfy that at least four species are consistently early in RT in more than 90% regions of this state, we assign this state to the weakly conserved early (WE) group. We assign states to the L and WL groups in a similar way accordingly. The remaining states are assigned to the NC group. Specifically, states 1-4 and states 5-8 are E states and L states, respectively. States 19-22, and states 13, 16, 23, 24 are WE and WL states, respectively. States 9-18 and 25-30 are NC states.

##### A.10 Alignment between the predicted RT states and TADs

When calculating the distances between the TAD boundaries and the RT state boundaries, to filter the TADs that are far away from any predicted states, we extended each boundary of a TAD with 30kb and used the states that overlap with the extended TAD to calculate the boundary distance. We then calculated the percentages of boundary distances that fall into four intervals. The first interval is [0,12kb]. The remaining three intervals are determined by the empirical distance distribution obtained from TAD shuffling, and equally cover the distances that are larger than 12kb. We shuffled the TADs 1000 times by randomly relocating them along the genome. We calculated and merged the boundary distances of each shuffle of TADs to form the empirical boundary distance distribution. Furthermore, for each shuffle of TADs, we computed the percentage of boundary distances that fall into each distance interval to form empirical distributions for each interval.

##### A.11 Motif feature analysis in lineage-specific RT states predicted by Phylo-HMGP

We performed motif scanning in the orthologous open chromatin regions of each of the five primate species. We identified open chromatin regions in human genome as DNase-seq peak regions with +/-250bp extension, using DNase-seq data of the GM12878 cells in human downloaded from the ENCODE annotation data in the UCSC genome browser (Rosenbloom et al., 2012). We used the liftOver tool (Hinrichs et al., 2006) to project the identified open chromatin regions in human genome to genomes of the other primate species, obtaining orthologous open chromatin regions in other species. We used FIMO (Grant et al., 2011) and 635 position weight matrices (PWMs) of TF binding motifs from the JASPAR 2016 core vertebrate motif database (Mathelier et al., 2016) to perform motif scanning in the orthologous open chromatin regions of each species. In each orthologous region, we computed the motif frequency for each PWM within the open chromatin area for each species ($p$-value$<$1e-04 required for each motif). We then normalized the frequency by the open chromatin area size within this orthologous region.

To identify TF binding motifs that may be lineage-specifically enriched in predicted lineage-specific RT states, we used two types of tests and selected motifs that can pass both tests. First, within each lineage-specific RT state, we performed binomial tests to find the motifs that are significantly more enriched in the RT-specific species than expected ($p$-value$<$0.05). Second, for a motif that passes the binomial test in a lineage-specific RT state, we calculated the fold change of its motif frequency within the RT-specific species compared to the other species. To estimate the empirical $p$-value, we randomly sampled the same number of regions as the lineage-specific RT state from the whole genome for 2000 times, and calculated the same type of fold change to form empirical distribution. We selected the motifs that have empirical $p$-values$<$0.05.

### B Supplementary Results

#### B.1 Evaluation of Phylo-HMGP model in comparsion with other methods on RT data

We also compared Phylo-HMGP-OU with Gaussian-HMM method, GMM method, K-means clustering method, and Phylo-HMGP-BM on the Repli-seq dataset, based on the average performance from 10 repeated runs of each method. We have applied each method to the RT data for state prediction, with state number set to be 30, as estimated in Supplementary Methods A.8. However, there is no available ground truth for the RT data. For evaluation purpose, we constructed an evaluation state set. Specifically, we discretized the signals of each species into 5 levels, and identified 12 possible selected representative states (10 possible lineage-specific states and two conserved states) from all the combinations of the 5 levels in orthologous regions across the species. We fit a Gaussian-HMM with five hidden states independently for each species, with each state representing a discretized level of the RT signal values. High signal values (level 1 and 2) and low signal values (level 3 and 4) correspond to early phase and late phase in RT, respectively. For example, human early state represents early RT only in human and non-early RT in the other four species at the orthologous regions. The 10 possible lineage-specific states identified from discrete levels of RT signals are human early/late, chimpanzee early/late, orangutan early/late, gibbon early/late, and green monkey early/late, respectively. The two identified conserved states are conserved RT early and late, respectively. The 12 selected states cover around 60% of all the orthologous regions with cross-species RT signals. We constrained the evaluation of different methods to the regions where the 12 selected states are present. Within these regions, we used the 12 selected states as a known partition, and evaluated the relevance of the prediction of each method to this partition, using the evaluation metrics AMI, NMI, RI, and $F_1$ (Fig. S7). As each method predicted 30 states, which is a finer partition than 12 states, the evaluation measures are generally lower than those in the simulation studies. For example, regions in one state in the 12-state partition may be predicted to be in different states in the 30-state partition, which affects the $F_1$ score by reducing the Recall and also affects the other metrics. Also, the selected states used for comparison were estimated using predictions from Gaussian-HMM in each species, which would favor Gaussian-HMM and GMM. The regions where 12 selected states are present are less continuous than the whole genome regions and have weaker spatial dependence between regions, which again would favor GMM. However, Phylo-HMGP-OU still outperforms the other methods in each of the four evaluation metrics. Phylo-HMGP-BM ranks second in performance. Even though the 12-state partition is not exactly ground truth, it nevertheless demonstrates that Phylo-HMGP outperforms the other methods in the RT data application, which is consistent with the results from the simulation studies.

#### B.2 Evaluation based on *cis*-regulatory module evolution

In addition to the real data application on the Repli-seq data, we also applied the models to predict different states of *cis*-regulatory module (CRM) evolution along the genome using features only from DNA sequences. We focused on a recent dataset for promoters and enhancers marked by H3K4me3 and H3K27ac in vertebrate liver cells (Villar et al., 2015). We used four species, including human (hg19), macaque (rheMac2), marmoset (calJac3), and mouse (mm10). We used hg19 as the reference and divide it into 5 kb bins. For each of the orthologous regions, we used the method Cluster-Buster (Frith et al., 2003) to compute a CRM score for presence of homotypic motif clusters within this region of the respective species, using a selected collection of 382 position weight matrices of TF binding motifs from the JASPAR 2016 core vertebrate motif database (Mathelier et al., 2016). We only used expressed TFs in liver cell based on gene expression data of human liver from GSE61260 (Horvath et al., 2014). We computed CRM sores for the 286,287 orthologous regions across the four species. We applied Phylo-HMGP to perform state prediction along the genome, with the state number set to be 16.

Here we assumed that the calculated CRM scores are associated with the activities of regulatory elements (e.g., enhancers or promoters). The ChIP-seq data, which can be used to identify and validate the existence of regulatory elements such as enhancers or promoters, were used to prepare benchmarks to evaluate the performance of the proposed model Phylo-HMGP in discovering different CRM patterns across species.

We used the peak regions called from ChIP-seq data of histone modification H3K27ac and H3K4me3 (Villar et al., 2015) to evaluate the different states estimated by Phylo-HMGP. For enhancer-evolution associated state prediction, we segmented the reference genome into different benchmark states based on the species-specific distribution of H3K27ac peaks. We then compared the states predicted by Phylo-HMGP-OU with the benchmark states, in comparison with the results from Gaussian-HMM, K-means clustering, and Phylo-HMGP-BM. We also performed the state evaluation using the H3K4me3 dataset. The results are shown in Fig. S9. Phylo-HMGP-OU achieved the highest RI and $F_1$ score among the different methods in the four experiments. Although the overall accuracy of using the CRM score for predicting enhancer/promoter activities seems not high and it remains an open problem to more accurately predict regulatory region activities from genome sequence, our evaluation again demonstrates the general utility and advantage of Phylo-HMGP.

#### C Supplementary Tables

| Simulation | Method | AMI | NMI | ARI | Precision | Recall | $F_1$ |
|---|---|---|---|---|---|---|---|
| Dataset I-1 | Gaussian-HMM | 0.7863 | 0.8216 | 0.5941 | 0.8206 | 0.5569 | 0.6629 |
| Dataset I-1 | GMM | 0.7562 | 0.7985 | 0.6340 | **0.8903** | 0.5679 | 0.6932 |
| Dataset I-1 | Clustering | 0.6471 | 0.6926 | 0.4752 | 0.7539 | 0.4412 | 0.5558 |
| Dataset I-1 | Phylo-HMM-BM | 0.8190 | 0.8494 | 0.7801 | 0.8496 | 0.8122 | 0.8226 |
| Dataset I-1 | Phylo-HMM-OU ($\lambda_0 = 0$) | 0.7966 | 0.8183 | 0.6088 | 0.7973 | 0.5918 | 0.6786 |
| Dataset I-1 | Phylo-HMM-OU ($\lambda_0 = 4.0$) | **0.8309** | **0.8733** | **0.8405** | 0.8568 | **0.8986** | **0.8728** |
| Dataset I-2 | Gaussian-HMM | 0.5342 | 0.6901 | 0.2931 | 0.9666 | 0.2948 | 0.4519 |
| Dataset I-2 | GMM | 0.3446 | 0.4508 | 0.1884 | 0.8114 | 0.2335 | 0.3625 |
| Dataset I-2 | Clustering | 0.2692 | 0.3599 | 0.1157 | 0.7173 | 0.1768 | 0.2837 |
| Dataset I-2 | Phylo-HMM-BM | 0.5208 | 0.6621 | 0.3150 | 0.9531 | 0.3218 | 0.4810 |
| Dataset I-2 | Phylo-HMM-OU ($\lambda_0 = 0$) | 0.5315 | 0.6782 | 0.3149 | 0.9602 | 0.3191 | 0.4790 |
| Dataset I-2 | Phylo-HMM-OU ($\lambda_0 = 4.0$) | **0.7691** | **0.8229** | **0.7638** | **0.9811** | **0.7718** | **0.8565** |
| Dataset I-3 | Gaussian-HMM | 0.7323 | 0.7912 | 0.4919 | 0.8314 | 0.4476 | 0.5819 |
| Dataset I-3 | GMM | 0.5536 | 0.6200 | 0.3354 | 0.7442 | 0.3057 | 0.4334 |
| Dataset I-3 | Clustering | 0.4973 | 0.5561 | 0.2798 | 0.6580 | 0.2741 | 0.3870 |
| Dataset I-3 | Phylo-HMM-BM | 0.6752 | 0.7277 | 0.4282 | 0.7646 | 0.4045 | 0.5283 |
| Dataset I-3 | Phylo-HMM-OU ($\lambda_0 = 0$) | 0.7146 | 0.7698 | 0.4741 | 0.8088 | 0.4372 | 0.5674 |
| Dataset I-3 | Phylo-HMM-OU ($\lambda_0 = 4.0$) | **0.8309** | **0.8733** | **0.8405** | **0.8568** | **0.8986** | **0.8728** |
| Dataset I-4 | Gaussian-HMM | 0.6965 | 0.7967 | 0.4824 | 0.9585 | 0.4482 | 0.6095 |
| Dataset I-4 | GMM | 0.7185 | 0.8106 | 0.5241 | 0.9737 | 0.4849 | 0.6457 |
| Dataset I-4 | Clustering | 0.5812 | 0.6810 | 0.3535 | 0.8928 | 0.3395 | 0.4911 |
| Dataset I-4 | Phylo-HMM-BM | 0.8151 | 0.8644 | 0.7342 | **0.9756** | 0.7112 | 0.8174 |
| Dataset I-4 | Phylo-HMM-OU ($\lambda_0 = 0$) | 0.7946 | 0.8544 | 0.6846 | 0.9736 | 0.6589 | 0.7776 |
| Dataset I-4 | Phylo-HMM-OU ($\lambda_0 = 4.0$) | **0.8396** | **0.8607** | **0.8146** | 0.9687 | **0.8071** | **0.8792** |
| Dataset I-5 | Gaussian-HMM | 0.7688 | 0.8331 | 0.5644 | 0.9091 | 0.4908 | 0.6374 |
| Dataset I-5 | GMM | 0.6398 | 0.6949 | 0.5065 | 0.8366 | 0.4552 | 0.5893 |
| Dataset I-5 | Clustering | 0.5298 | 0.5906 | 0.3494 | 0.7227 | 0.3218 | 0.4453 |
| Dataset I-5 | Phylo-HMM-BM | 0.7654 | 0.8243 | 0.5594 | 0.8908 | 0.4929 | 0.6346 |
| Dataset I-5 | Phylo-HMM-OU ($\lambda_0 = 0$) | 0.7738 | 0.8370 | 0.5809 | 0.9180 | 0.5057 | 0.6521 |
| Dataset I-5 | Phylo-HMM-OU ($\lambda_0 = 4.0$) | **0.8798** | **0.9187** | **0.9594** | **0.9524** | **0.9867** | **0.9692** |
| Dataset I-6 | Gaussian-HMM | 0.8736 | **0.9141** | 0.7466 | **0.9642** | 0.6635 | 0.7861 |
| Dataset I-6 | GMM | 0.7810 | 0.8157 | 0.6810 | 0.8884 | 0.6215 | 0.7312 |
| Dataset I-6 | Clustering | 0.6409 | 0.6785 | 0.5101 | 0.7457 | 0.4791 | 0.5833 |
| Dataset I-6 | Phylo-HMM-BM | 0.8554 | 0.8772 | 0.7534 | 0.9052 | 0.7133 | 0.7932 |
| Dataset I-6 | Phylo-HMM-OU ($\lambda_0 = 0$) | 0.8538 | 0.8891 | 0.7216 | 0.9325 | 0.6490 | 0.7652 |
| Dataset I-6 | Phylo-HMM-OU ($\lambda_0 = 4.0$) | **0.8906** | 0.8958 | **0.8390** | 0.9224 | **0.8200** | **0.8678** |

**Table S1:** Related to Fig. 2. Performance evaluation of Gaussian-HMM, GMM (Gaussian Mixture Model), K-means Clustering, Phylo-HMGP-BM, Phylo-HMGP-OU ($\lambda_0 = 0$), and Phylo-HMGP-OU ($\lambda_0 = 4.0$) on six simulated datasets in Simulation Study I with respect to AMI (Adjusted Mutual Information), NMI (Normalized Mutual Information), ARI (Adjusted Rand Index), Precision, Recall, and $F_1$ score. Each method is repeated 10 times with different initializations on each simulation dataset. The average performance from the 10 repeated runs of each method is presented. The best performance of the compared methods is in bold font.

| Simulation | Method | AMI | NMI | ARI | Precision | Recall | $F_1$ |
|---|---|---|---|---|---|---|---|
| Dataset II-1 | Gaussian-HMM | 0.8314 | 0.8765 | 0.6693 | **0.9244** | 0.5965 | 0.7251 |
| Dataset II-1 | GMM | 0.7399 | 0.7823 | 0.6138 | 0.8800 | 0.5511 | 0.6776 |
| Dataset II-1 | Clustering | 0.6144 | 0.6623 | 0.4509 | 0.7579 | 0.4124 | 0.5341 |
| Dataset II-1 | Phylo-HMM-BM | 0.8511 | **0.8796** | **0.7786** | 0.8785 | **0.7842** | **0.8205** |
| Dataset II-1 | Phylo-HMM-OU | **0.8534** | 0.8706 | 0.7385 | 0.8319 | 0.7591 | 0.7906 |
| Dataset II-2 | Gaussian-HMM | 0.5328 | 0.6863 | 0.2957 | 0.9550 | 0.3099 | 0.4680 |
| Dataset II-2 | GMM | 0.3421 | 0.4477 | 0.1893 | 0.8206 | 0.2416 | 0.3733 |
| Dataset II-2 | Clustering | 0.2590 | 0.3481 | 0.1119 | 0.7278 | 0.1780 | 0.2860 |
| Dataset II-2 | Phylo-HMM-BM | 0.5343 | 0.6877 | 0.2944 | 0.9550 | 0.3085 | 0.4663 |
| Dataset II-2 | Phylo-HMM-OU | **0.6901** | **0.7833** | **0.5883** | **0.9736** | **0.6033** | **0.7353** |
| Dataset II-3 | Gaussian-HMM | 0.7754 | 0.8307 | 0.5236 | **0.8469** | 0.4700 | 0.6045 |
| Dataset II-3 | GMM | 0.5641 | 0.6248 | 0.3463 | 0.7225 | 0.3176 | 0.4412 |
| Dataset II-3 | Clustering | 0.5098 | 0.5606 | 0.2991 | 0.6385 | 0.2963 | 0.4048 |
| Dataset II-3 | Phylo-HMM-BM | 0.7239 | 0.7727 | 0.4652 | 0.7793 | 0.4338 | 0.5561 |
| Dataset II-3 | Phylo-HMM-OU | **0.8159** | **0.8952** | **0.8797** | **0.8469** | **0.9848** | **0.9105** |
| Dataset II-4 | Gaussian-HMM | 0.6585 | 0.7623 | 0.4139 | 0.9318 | 0.3898 | 0.5497 |
| Dataset II-4 | GMM | 0.7451 | 0.8262 | 0.5859 | **0.9778** | 0.5486 | 0.7027 |
| Dataset II-4 | Clustering | 0.5411 | 0.6463 | 0.2933 | 0.8678 | 0.2868 | 0.4307 |
| Dataset II-4 | Phylo-HMM-BM | 0.7857 | 0.8411 | 0.6904 | 0.9543 | 0.6812 | 0.7775 |
| Dataset II-4 | Phylo-HMM-OU | **0.8092** | **0.8556** | **0.7135** | 0.9583 | **0.7045** | **0.7981** |
| Dataset II-5 | Gaussian-HMM | 0.7932 | 0.8529 | 0.5842 | 0.9078 | 0.5125 | 0.6551 |
| Dataset II-5 | GMM | 0.6640 | 0.7155 | 0.5399 | 0.8595 | 0.4827 | 0.6173 |
| Dataset II-5 | Clustering | 0.5317 | 0.5894 | 0.3375 | 0.7028 | 0.3142 | 0.4343 |
| Dataset II-5 | Phylo-HMM-BM | 0.7999 | 0.8595 | 0.5891 | 0.9223 | 0.5139 | 0.6573 |
| Dataset II-5 | Phylo-HMM-OU | **0.9162** | **0.9454** | **0.9504** | **0.9555** | **0.9694** | **0.9622** |
| Dataset II-6 | Gaussian-HMM | 0.8600 | 0.8969 | 0.7162 | **0.9411** | 0.6396 | 0.7613 |
| Dataset II-6 | GMM | 0.7742 | 0.8118 | 0.6665 | 0.8887 | 0.6050 | 0.7197 |
| Dataset II-6 | Clustering | 0.6319 | 0.6733 | 0.5036 | 0.7554 | 0.4689 | 0.5784 |
| Dataset II-6 | Phylo-HMM-BM | 0.8371 | 0.8612 | 0.7241 | 0.8788 | 0.6944 | 0.7701 |
| Dataset II-6 | Phylo-HMM-OU | **0.9117** | **0.9150** | **0.8632** | 0.9314 | **0.8506** | **0.8888** |

**Table S2:** Related to Fig. 2. Performance evaluation of Gaussian-HMM, GMM, K-means Clustering, Phylo-HMGP-BM, and Phylo-HMGP-OU ($\lambda_0 = 4.0$) on six simulated datasets in Simulation Study II with respect to AMI, NMI, ARI, Precision, Recall, and $F_1$ score. Each method is repeated 10 times with different initializations on each simulation dataset. The average performance from the 10 repeated runs of each method is presented. The best performance of the compared methods is in bold font.

| Conserved RT early | GO term/Pathway | Count | Fold enrichment | $p$-value |
|---|---|---|---|---|
| Constitutive RT early | intracellular transport | 431 | 1.2 | 6.4e-06 |
| | amide biosynthetic process | 214 | 1.3 | 1.5e-05 |
| | posttranscriptional regulation of gene expression | 149 | 1.4 | 2.0e-05 |
| | cellular amide metabolic process | 277 | 1.2 | 2.0e-05 |
| | peptide biosynthetic process | 195 | 1.3 | 2.4e-05 |
| | peptide metabolic process | 231 | 1.3 | 2.5e-05 |
| | mRNA metabolic process | 193 | 1.2 | 5.8e-05 |
| | translation | 187 | 1.3 | 5.9e-05 |
| | microtubule-based process | 179 | 1.3 | 1.0e-04 |
| | clathrin-mediated endocytosis | 22 | 2.1 | 3.0e-04 |
| | regulation of vascular permeability | 18 | 2.3 | 3.9e-04 |
| | mitochondrion organization | 183 | 1.2 | 7.2e-04 |
| Non-constitutive RT early | immune response | 540 | 1.3 | 7.9e-013 |
| | regulation of immune response | 327 | 1.3 | 2.7e-010 |
| | defense response | 520 | 1.2 | 5.5e-08 |
| | symbiosis, encompassing mutualism through parasitism | 376 | 1.2 | 5.7e-08 |
| | interspecies interaction between organisms | 376 | 1.2 | 5.7e-08 |
| | viral process | 362 | 1.2 | 1.2e-07 |
| | multi-organism cellular process | 364 | 1.2 | 1.7e-07 |
| | immune effector process | 257 | 1.3 | 2.6e-07 |
| | immune response-activating signal transduction | 176 | 1.4 | 7.9e-07 |
| | activation of immune response | 192 | 1.3 | 1.1e-06 |
| | immune response-regulating signaling pathway | 186 | 1.3 | 1.2e-06 |
| | regulation of immune system process | 465 | 1.2 | 2.5e-06 |

**Table S3:** Related to Fig. 3. Gene ontology (GO) terms or pathways that show significant correlation with regions that are both constitutive RT early and conserved RT early, and regions that are conserved RT early but not constitutive RT early. We used DAVID to perform the gene ontology analysis. The column Count represents the number of genes found to be associated with the corresponding GO term/pathway in the queried regions. The conserved early RT regions are based on state prediction by Phylo-HMGP. We observed that genes enriched in the regions that are both constitutive early and conserved early are mainly involved in basic biological functions and processes that are shared between different cell types. Genes associated with the regions that are not constitutive early but conserved early are involved in the cell type specific functions of the lymphoblastoid cells, such as the immune response functions.

| Predicted state | GO term/Pathway | Count | Fold enrichment | $p$-value |
|---|---|---|---|---|
| State 9 | sensory perception of smell | 16 | 8.5 | 1.9e-09 |
| | detection of chemical stimulus | 17 | 6.6 | 6.3e-09 |
| | detection of stimulus involved in sensory perception | 17 | 6.4 | 9.2e-09 |
| | olfactory transduction | 16 | 6.0 | 4.1e-08 |
| | sensory perception | 21 | 2.9 | 3.6e-05 |
| | neurological system process | 23 | 2.2 | 8.2e-04 |
| State 10 | regulation of nucleic acid-templated transcription | 95 | 1.5 | 1.4e-05 |
| | regulation of RNA biosynthetic process | 95 | 1.5 | 1.9e-05 |
| | regulation of nucleobase-containing compound metabolic process | 101 | 1.4 | 1.4e-04 |
| | aromatic compound biosynthetic process | 107 | 1.4 | 1.5e-04 |
| | heterocycle biosynthetic process | 106 | 1.4 | 2.1e-04 |
| | regulation of nitrogen compound metabolic process | 104 | 1.3 | 5.8e-04 |
| | cellular macromolecule biosynthetic process | 114 | 1.3 | 1.1e-03 |
| State 11 | G-protein coupled purinergic nucleotide receptor signaling pathway | 4 | 33.2 | 2.0e-04 |
| | cellular response to organic substance | 32 | 1.8 | 1.6e-03 |
| | insulin secretion | 6 | 5.0 | 6.7e-03 |
| | response to endogenous stimulus | 23 | 1.8 | 9.6e-03 |
| State 14 | endocardial cushion morphogenesis | 7 | 15.6 | 4.4e-06 |
| | cartilage development | 13 | 4.9 | 1.4e-05 |
| | endocardial cushion development | 7 | 11.6 | 2.7e-05 |
| | connective tissue development | 14 | 4.1 | 4.1e-05 |
| | pattern specification process | 18 | 2.8 | 2.1e-04 |
| | organ morphogenesis | 29 | 2.0 | 5.1e-04 |
| | enzyme linked receptor protein signaling pathway | 27 | 2.0 | 1.1e-03 |
| | circulatory system development | 27 | 2.0 | 1.3e-03 |
| | cardiovascular system development | 27 | 2.0 | 1.3e-03 |
| State 16 | cell adhesion | 27 | 2.2 | 1.4e-04 |
| | regulation of nervous system development | 15 | 2.6 | 1.5e-03 |
| | regulation of neurogenesis | 13 | 2.6 | 4.3e-03 |
| | regulation of cellular component organization | 29 | 1.7 | 4.5e-03 |
| | neuron projection morphogenesis | 11 | 2.8 | 6.1e-03 |
| | neuron differentiation | 18 | 2.0 | 8.3e-03 |
| State 18 | regulation of locomotion | 32 | 2.6 | 1.4e-06 |
| | regulation of cell motility | 31 | 2.7 | 1.6e-06 |
| | regulation of cell migration | 29 | 2.7 | 3.5e-06 |
| | regulation of cell differentiation | 45 | 1.9 | 4.0e-05 |
| | epithelium development | 33 | 1.9 | 3.4e-04 |
| | cell proliferation | 43 | 1.5 | 6.6e-03 |
| | regulation of anatomical structure morphogenesis | 28 | 1.7 | 7.8e-03 |
| | movement of cell or subcellular component | 42 | 1.5 | 8.3e-03 |

**Table S4:** Related to Fig. 4. Example gene ontology (GO) terms or pathways that show significant correlation with lineage-specific RT states (State 9: human-chimpanzee specific early RT state; State 10: human-chimpanzee specific late RT state; State 11: human-chimpanzee-orangutan specific early RT state; State 14: orangutan specific late RT state; State 16: gibbon specific late RT state; State 18: green monkey specific late RT state). We use DAVID to perform the gene ontology analysis.

#### D Supplementary Figures

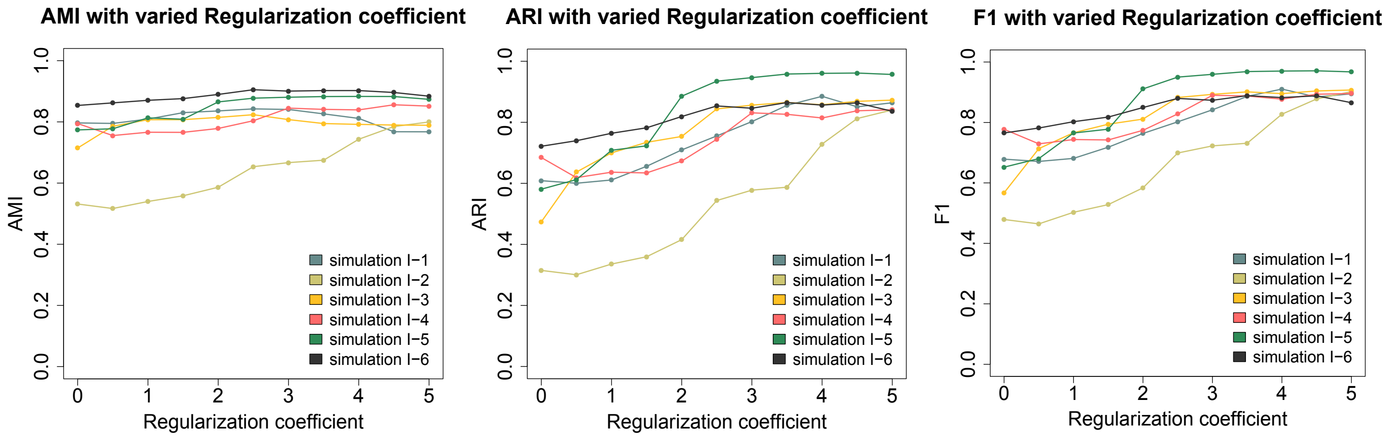


**Figure S1:** Related to Fig. 2. Performance evaluation on AMI, ARI, and $F_1$ score in Simulation Study I with respect to varied $l_2$-norm regularization coefficient $\lambda_0$.

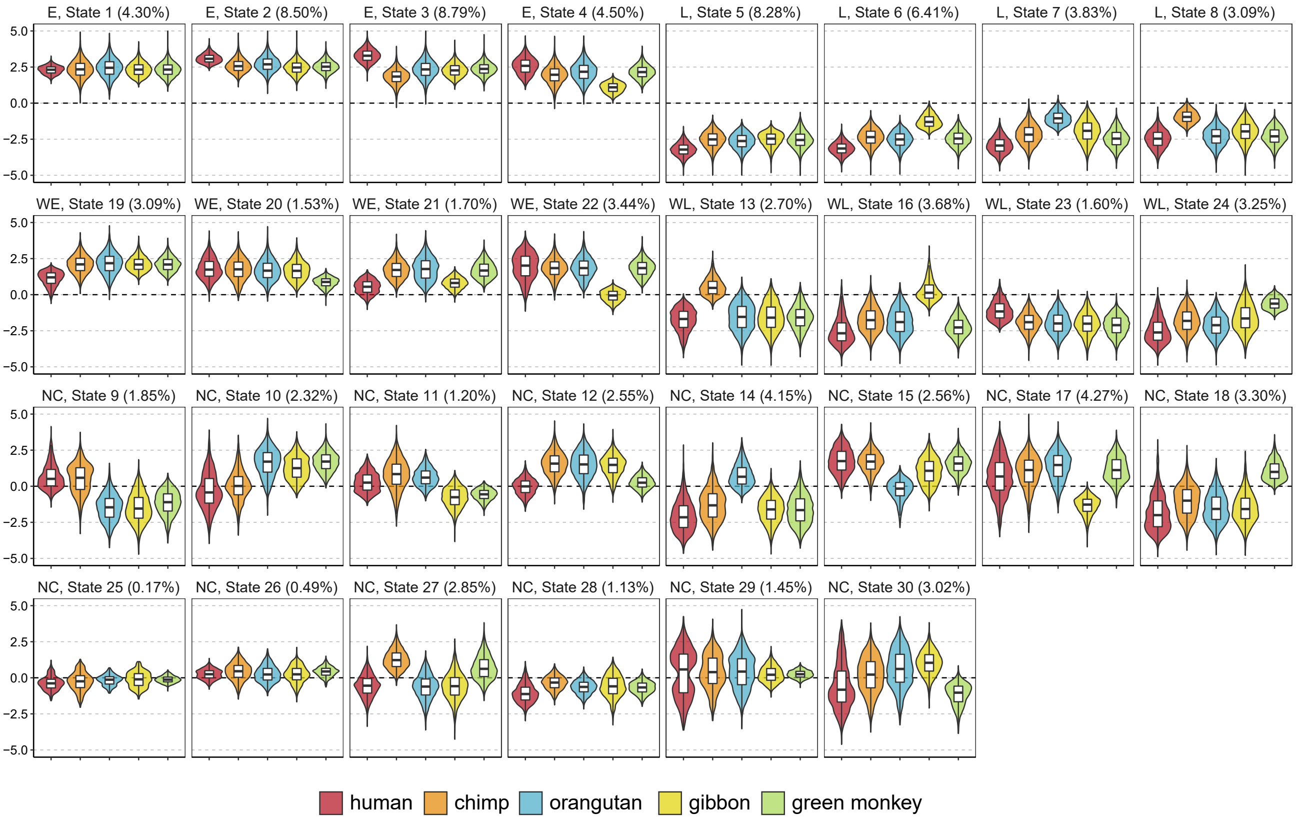


**Figure S2:** Related to Fig. 3. Different patterns of replication timing (RT) across five primate species predicted by Phylo-HMGP-OU for 30 states. The y-axis represents the RT signal value. Box plots of the RT signal distributions of the five primate species in each predicted state are shown. The percentage of the number of regions in each predicted state and the RT group label are also shown in the title of each plot.

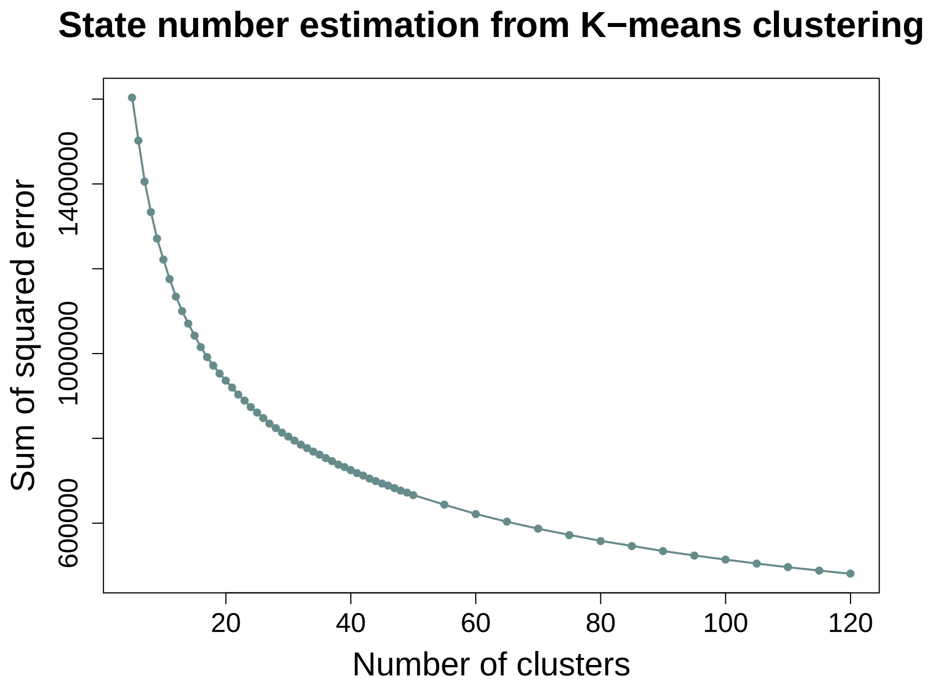


**Figure S3:** Related to Fig. 3. The change of Sum of Squared Error (SSE) with respect to an increased cluster number in $K$-means clustering on RT data. The state number was estimated to be between 20 and 40 based on the results from $K$-means clustering.

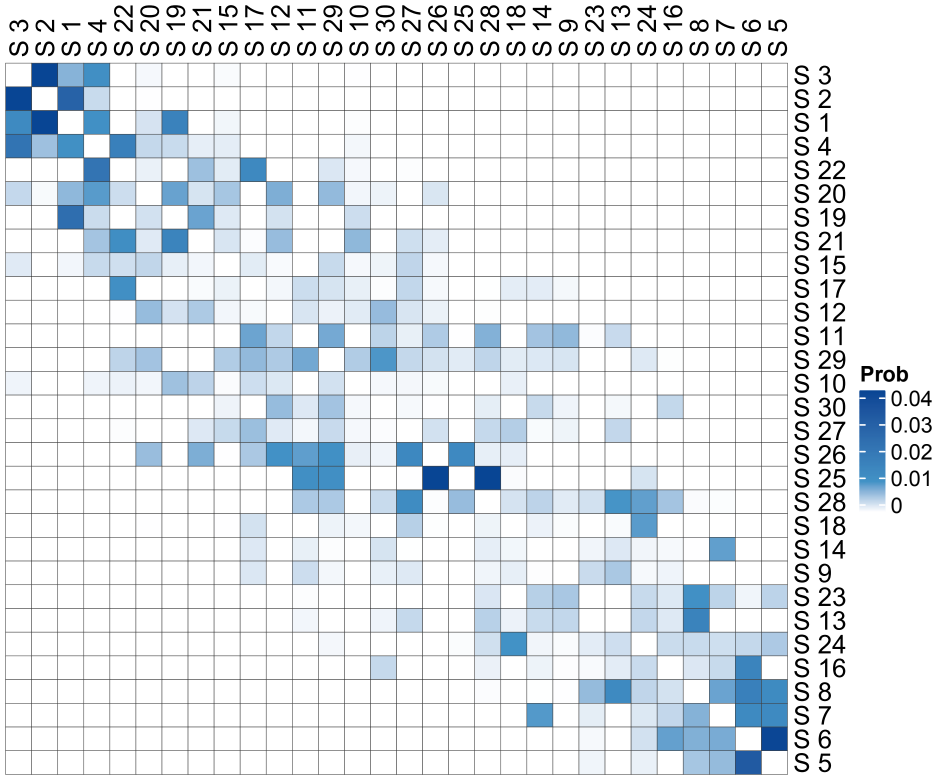


**Figure S4:** Related to Fig. 3. Transition probability matrix of the 30 states estimated by Phylo-HMGP-OU. The self-transition probability is not shown (set to be blank) to illustrate the probabilities of transitions to other states more significantly.

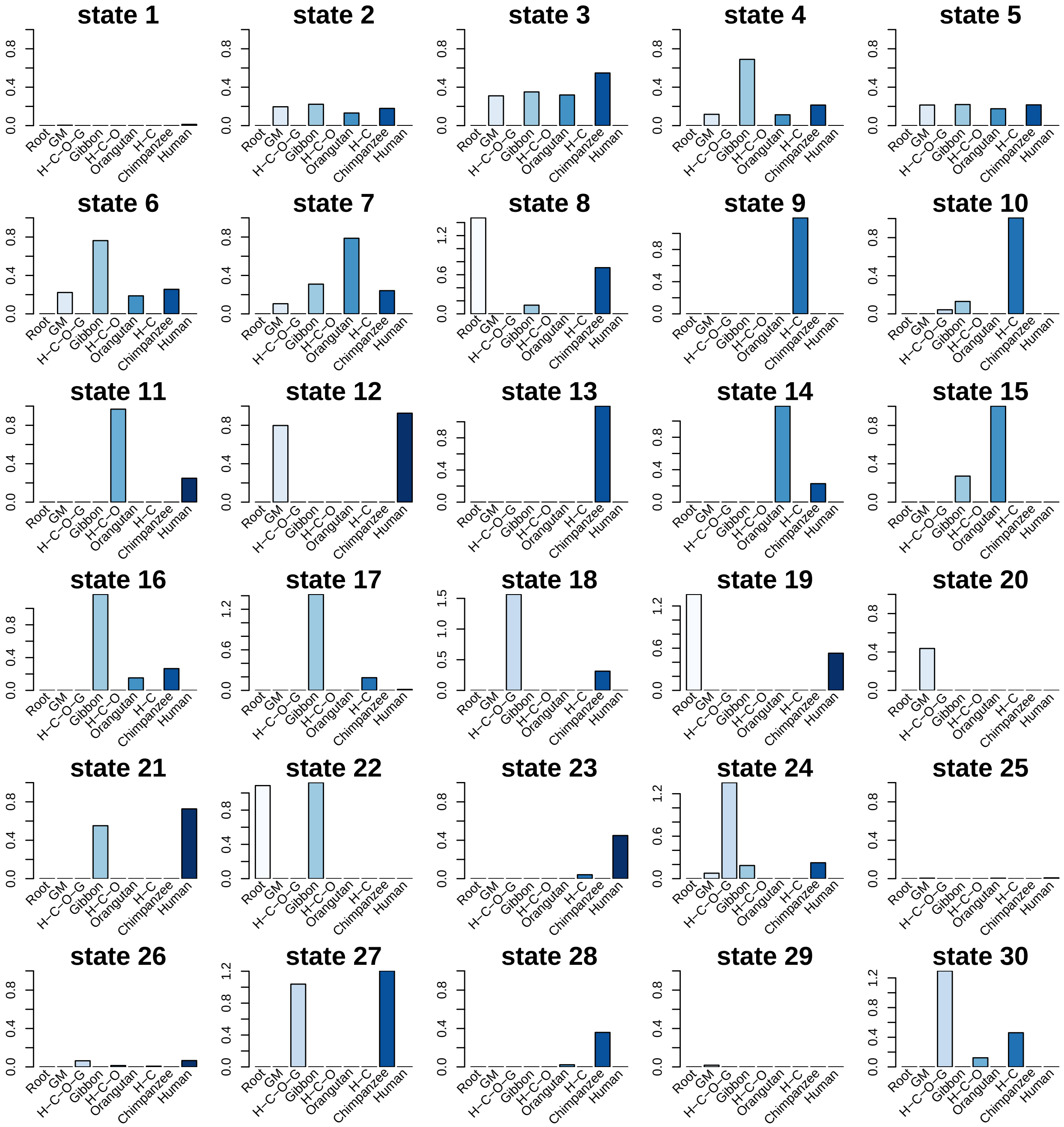

**Figure S5:** Related to Fig. 3. Estimated Brownian motion intensity along each branch of the phylogenetic tree in the 30 states predicted by Phylo-HMGP-OU. 'GM' stands for Green Monkey. Each column corresponds to the branch connecting the nearest ancestor of the species specified by the species name to the species. Root stands for the branch connecting the remote root node ancestor with the nearest common ancestor of green monkey and human. H-C-O-G, H-C-O, and H-C represent the branches leading to the clade of human, chimpanzee, orangutan, and gibbon, the clade of human, chimpanzee, and orangutan, and the clade of human and chimpanzee, respectively.

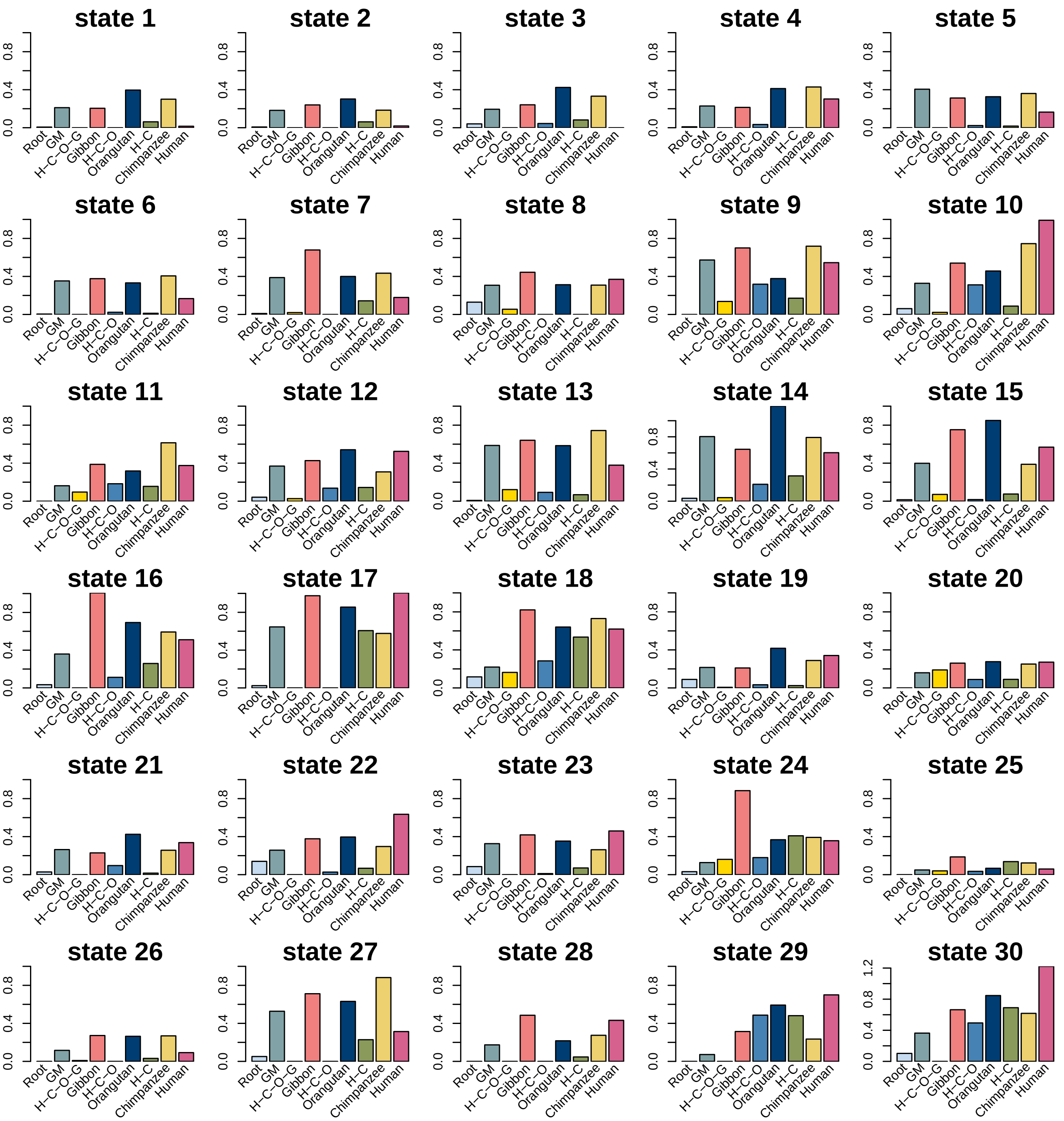


**Figure S6:** Related to Fig. 3. Estimated selection strength along each branch of the phylogenetic tree in the 30 states predicted by Phylo-HMGP-OU. 'GM' stands for Green Monkey. Each column corresponds to the branch connecting the nearest ancestor of the species specified by the species name to the species. Root stands for the branch connecting the remote root node ancestor with the nearest common ancestor of green monkey and human. H-C-O-G, H-C-O, and H-C represent the branches leading to the clade of human, chimpanzee, orangutan, and gibbon, the clade of human, chimpanzee, and orangutan, and the clade of human and chimpanzee, respectively.

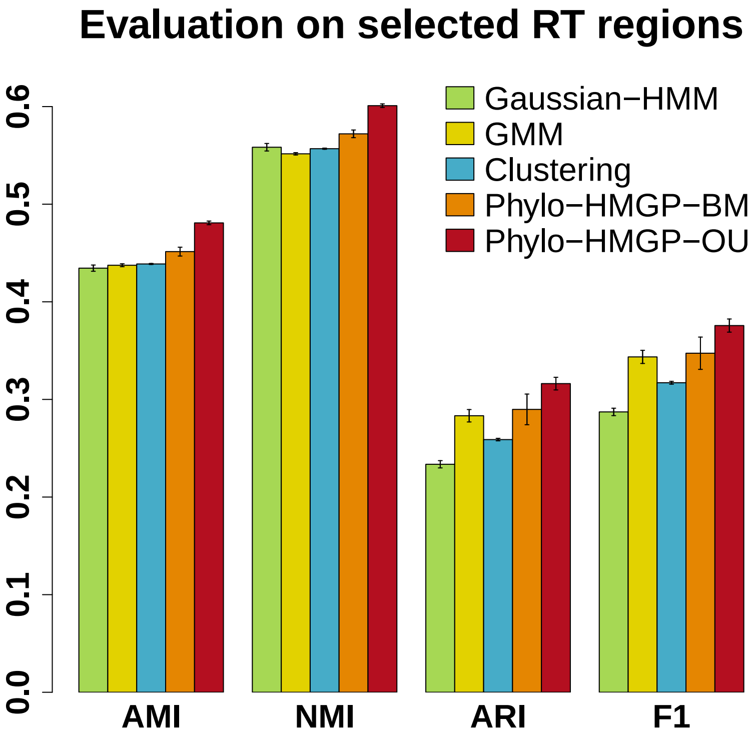


**Figure S7:** Related to Fig. 3. Evaluation of Gaussian-HMM, GMM, K-means Clustering, Phylo-HMGP-BM, and Phylo-HMGP-OU on the replication timing dataset in terms of AMI, NMI, ARI and $F_1$ score in the genomic regions of 12 different states (including 10 lineage-specific states and two conserved states) identified from comparison of discretized single-species observations. The standard error of the results of 10 repeated runs for each method is shown as the error bar.

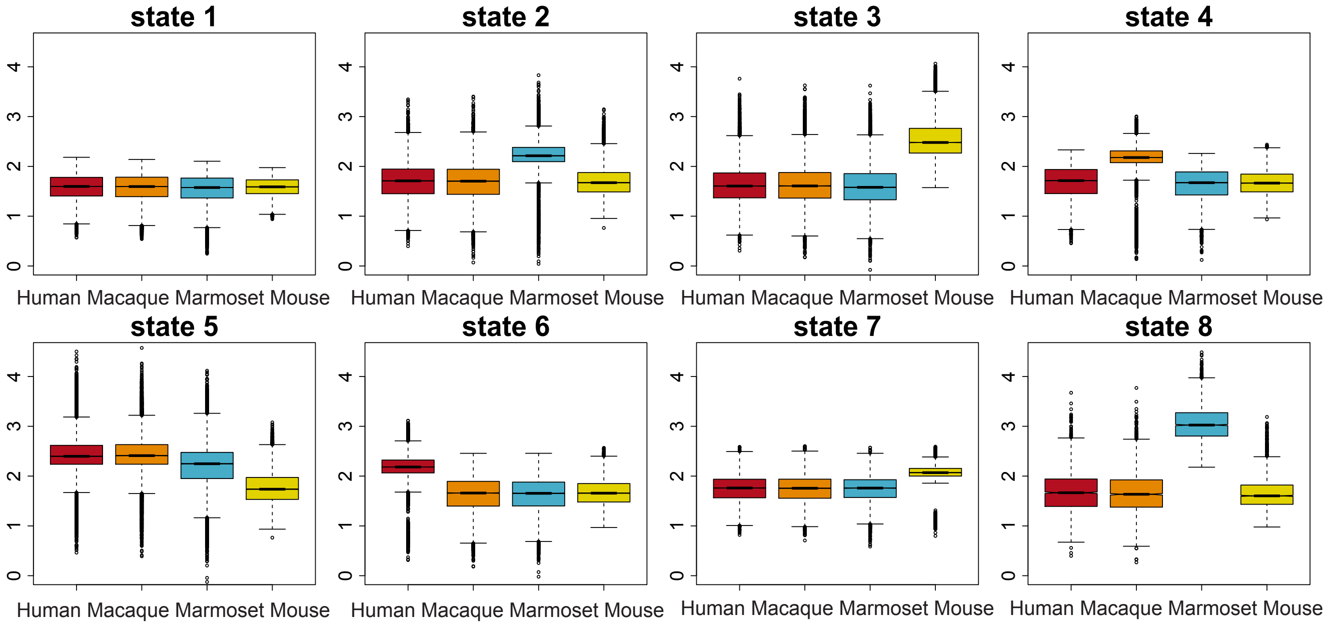


**Figure S8:** Related to Fig. 1. Different patterns of CRM (*cis*-regulatory module) score across four mammalian species (human, macaque, marmoset, and mouse) predicted by Phylo-HMGP-OU. Box plots of the top 8 states with the most number of genomic regions are shown. The y-axis represents the logarithm of the CRM score.

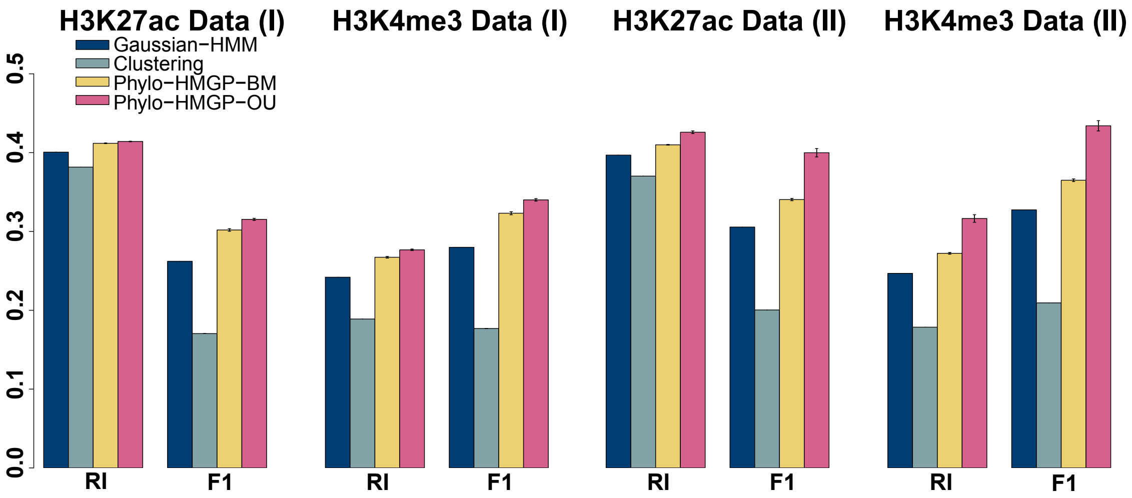


**Figure S9:** Related to Fig. 1. Evaluation of Gaussian-HMM, K-means Clustering, Phylo-HMGP-BM, and Phylo-HMGP-OU on H3K27ac and H3K4me3 ChIP-seq datasets in terms of RI (Rand Index), and $F_1$ score. Experiments (I) represent they are perforemd for the four mammal species. Experiments (II) represent they are performed for the three primate species human, macaque, and marmoset. The standard error of the results of 10 repeated runs for each method is shown as the error bar.

#### References


J. A. Bilmes et al. A gentle tutorial of the em algorithm and its application to parameter estimation for gaussian mixture and hidden markov models. 1998.

D. L. Davies and D. W. Bouldin. A cluster separation measure. *IEEE transactions on pattern analysis and machine intelligence*, (2):224–227, 1979.

N. Day, A. Hemmaplardh, R. E. Thurman, J. A. Stamatoyannopoulos, and W. S. Noble. Unsupervised segmentation of continuous genomic data. *Bioinformatics*, 23(11):1424–1426, 2007.

M. C. Frith, M. C. Li, and Z. Weng. Cluster-buster: Finding dense clusters of motifs in dna sequences. *Nucleic acids research*, 31(13):3666–3668, 2003.

C. E. Grant, T. L. Bailey, and W. S. Noble. Fimo: scanning for occurrences of a given motif. *Bioinformatics*, 27(7):1017–1018, 2011.

A. S. Hinrichs, D. Karolchik, R. Baertsch, G. P. Barber, G. Bejerano, H. Clawson, M. Diekhans, T. S. Furey, R. A. Harte, F. Hsu, et al. The ucsc genome browser database: update 2006. *Nucleic acids research*, 34(suppl_1):D590–D598, 2006.

S. Horvath, W. Erhart, M. Brosch, O. Ammerpohl, W. von Schönfels, M. Ahrens, N. Heits, J. T. Bell, P.-C. Tsai, T. D. Spector, et al. Obesity accelerates epigenetic aging of human liver. *Proceedings of the National Academy of Sciences*, 111(43):15538–15543, 2014.

C. D. Manning, P. Raghavan, and H. Schütze. *Introduction to Information Retrieval*. Cambridge University Press, New York, NY, USA, 2008. ISBN 0521865719, 9780521865715.

A. Mathelier, O. Fornes, D. J. Arenillas, C.-y. Chen, G. Denay, J. Lee, W. Shi, C. Shyr, G. Tan, R. Worsley-Hunt, et al. Jaspar 2016: a major expansion and update of the open-access database of transcription factor binding profiles. *Nucleic acids research*, 44(D1):D110–D115, 2016.

F. Pedregosa, G. Varoquaux, A. Gramfort, V. Michel, B. Thirion, O. Grisel, M. Blondel, P. Prettenhofer, R. Weiss, V. Dubourg, J. Vanderplas, A. Passos, D. Cournapeau, M. Brucher, M. Perrot, and E. Duchesnay. Scikit-learn: Machine learning in Python. *Journal of Machine Learning Research*, 12: 2825–2830, 2011.

D. B. Percival and A. T. Walden. *Wavelet methods for time series analysis*, volume 4. Cambridge university press, 2006.

B. D. Pope, T. Ryba, V. Dileep, F. Yue, W. Wu, O. Denas, D. L. Vera, Y. Wang, R. S. Hansen, T. K. Canfield, et al. Topologically associating domains are stable units of replication-timing regulation. *Nature*, 515(7527):402–405, 2014.

L. R. Rabiner. A tutorial on hidden markov models and selected applications in speech recognition. *Proceedings of the IEEE*, 77(2):257–286, 1989.

K. R. Rosenbloom, C. A. Sloan, V. S. Malladi, T. R. Dreszer, K. Learned, V. M. Kirkup, M. C. Wong, M. Maddren, R. Fang, S. G. Heitner, et al. Encode data in the ucsc genome browser: year 5 update. *Nucleic acids research*, 41(D1):D56–D63, 2012.

G. H. Thomas, R. P. Freckleton, and T. Székely. Comparative analyses of the influence of developmental mode on phenotypic diversification rates in shorebirds. *Proceedings of the Royal Society of London B: Biological Sciences*, 273(1594):1619–1624, 2006.

G. H. Thomas, S. Meiri, and A. B. Phillimore. Body size diversification in anolis: novel environment and island effects. *Evolution*, 63(8):2017–2030, 2009.

D. Villar, C. Berthelot, S. Aldridge, T. F. Rayner, M. Lukk, M. Pignatelli, T. J. Park, R. Deaville, J. T. Erichsen, A. J. Jasinska, et al. Enhancer evolution across 20 mammalian species. *Cell*, 160(3):554–566, 2015.

N. X. Vinh, J. Epps, and J. Bailey. Information theoretic measures for clusterings comparison: Variants, properties, normalization and correction for chance. *Journal of Machine Learning Research*, 11(Oct):

2837–2854, 2010.

P. Zwiernik, C. Uhler, and D. Richards. Maximum likelihood estimation for linear gaussian covariance models. *Journal of the Royal Statistical Society: Series B (Statistical Methodology)*, 79(4):1269–1292, 2017.
